# Ubiquitin Ligase SmDDA1b of Eggplant (*Solanum melongena*) Enhances Bacterial Wilt Resistance via SmNAC Degradation

**DOI:** 10.1101/2021.12.03.471130

**Authors:** Yixi Wang, Shuangshuang Yan, Bingwei Yu, Yuwei Gan, Jiangjun Lei, Changming Chen, Zhangsheng Zhu, Zhengkun Qiu, Bihao Cao

## Abstract

Bacterial wilt (BW) is a soil-borne disease that severely impacts plant growth and productivity globally. Ubiquitination plays a crucial role in disease resistance. Our previous research indicated that NAC transcription factor SmNAC negatively regulates BW resistance in eggplant (*Solanum melongena*). However, whether the ubiquitin/26S proteasome system (UPS) participates in this regulation is unknown.

This study used SmNAC as a bait to screen eggplant cDNA library and obtained SmDDA1b, an E3 ubiquitin ligase. Subcellular location and bimolecular fluorescence complementation assays revealed that SmDDA1b could interact with SmNAC in the nucleus. The *in vivo* and *in vitro* ubiquitination experiments indicated that SmDDA1b can degrade SmNAC through UPS. However, the discovery of negative regulation of SmDDA1b expression by SmNAC showed that there was a negative feedback loop between SmNAC and SmDDA1b in eggplant.

The *SmDDA1b-*overexpressed lines showed a higher BW resistance associated with high expression levels of salicylic acid (SA)-related genes and SA content than the wild-type lines. However, *SmDDA1b*-silencing lines showed the opposite results, indicating that *SmDDA1b* is a positive regulatory gene for BW resistance.

This study provides a candidate gene that can enhance BW resistance in eggplants. In addition, it provides insight into a mechanism that promotes plant disease resistance via the SmDDA1b-SmNAC-SA pathway.

## Introduction

Bacterial wilt (BW) is a soil-borne bacterial disease caused by *Ralstonia solanacearum* species complex (RSSC) (Safni *et al*., 2014), with diverse strains and a wide range of hosts. Hayward (1991) estimates that it can infect about 450 plant species in 54 families, including major cash crops and vegetables, especially *Solanaceae* crops. RSSC is one of the most common bacteria causing severe plant diseases globally (Mansfield et al., 2012; Kim et al., 2016). RSSC enters the plant xylem through the intercellular space for self-reproduction and secretes several extracellular polysaccharides and extracellular proteases. This blocks the vascular bundles and causing plants to die due to lack of water (McGarvey et al., 1999; Huang and Allen, 2000).

However, plants also resist BW in multiple ways, regulated at the DNA, transcription, translation, and post-translational levels. Studies have shown that heterologous overexpression of *Arabidopsis thaliana* gene *AtEFR* can reduce the effect of BW in tomato (*Solanum lycopersicum*) and potato (*Solanum tuberosum*) (Boschi *et al*., 2017; Kunwar *et al*., 2018). StNACb4 transcription factor positively regulates BW resistance in tomatoes at the transcriptional level (Chang *et al*., 2020). Moreover, transcription factor bHLH93 interacts with RSSC effector Ripl to induce plant immune response in tobacco (*Nicotiana tabacum*) (Tahir *et al*., 2017). RRS1-R can recognize RSSC Avr protein in *Arabidopsis* and act as dual resistance proteins with RPS4 for disease resistance (Tasset et al., 2010; Narusaka et al., 2014). Gong *et al*. (2021) showed that some histone deacetylase (HDAC)-mediated histone acetylation can reduce tomato resistance to BW at the post-translation level. Besides, Yu *et al*. (2020) found that phosphorylation of the *SGT1* gene is beneficial to BW resistance.

Isochorismate synthase (ICS) and phenylalanine ammonia lyase (PAL) synthesize salicylic acid (SA). Besides, the ICS pathway synthesizes more than 90% of SA in disease resistance response (Wildermuth et al., 2001; Garcion et al., 2008). There are two *ICS* genes in *Arabidopsis, ICS1*/*SID2* and *ICS2* (Dempsey *et al*., 2011). MdWRKY15 increases SA accumulation via *MdICS1* activation (Zhao *et al*., 2020). OsWRKY6 increases SA content via *OsICS1* activation (Choi *et al*., 2015). Lowe-Power *et al*. (2016) showed that SA can inhibit the expression of type III effectors of RSSC. External application of SA can increase *CaWRKY22* expression, thus enhancing BW resistance to pepper (*Capsicum annuum*) (Hussain *et al*., 2018). *NtWRKY50* overexpression enhances BW resistance in tobacco while significantly increasing SA levels (Liu *et al*., 2017). SA pathway signal genes also positively regulate plant disease resistance. Previous studies have shown that SA signaling transduction through *NPR, TGA, NPR1, TGA2*.*2*, and *TGA1a* positively regulates tomato BW resistance (Chen et al., 2009; Li et al., 2019). *EDS1, PAD4, NPR1* and *SGT1* positively regulates eggplant BW resistance (Xi-ou *et al*., 2016). *PR* gene expression is induced when the plant is stressed and the SA content increases (Lu *et al*., 2018).

Furthermore, ubiquitination is as important as phosphorylation and acetylation in eukaryotes. Ubiquitin/26S proteasome system (UPS) is a conserved ubiquitination system (Pickart and Fushman, 2004). Ubiquitin (Ub) interacts with the target protein in UPS through E1 (ubiquitin-activating) enzyme, E2 (ubiquitin-conjugating) enzyme, and E3 ubiquitin ligase via ATP for single ubiquitination or repeated polyubiquitination. This degrades or modifies the protein composition to regulate the function of eukaryotes (Thrower et al., 2000; Rowland et al., 2005). E3 ligases are mainly divided into HECT E3s, RING E3s, and RBR E3s. The RING family is the largest, containing a zinc or U-box binding domain (Stone et al., 2005; Morreale and Walden, 2016). Cullin-RING-Ligases (CRLs) are multi-subunit complexes and the largest family in the RING E3s. It is composed of scaffold protein Cullin, RBX1 protein-containing RING domain, adaptor, and substrate receptor (Zimmerman *et al*., 2010).

CUL1, CUL3, CUL4, and APC are the major cullin types in plants. CRL4 (CUL4A or CUL4B) uses DDB1 as an adaptor and DDB1 and cullin 4-related factors (DCFA) as substrate receptors (Pang *et al*., 2019). DDB1 and DET1-associated protein1 (DDA1) is a basic conservative component in the CRL4 core complex that directly interacts with DDB1 to promote substrate recruitment or regulate the overall topology of the CRL4-substrate complex (Olma *et al*., 2009; Shabek *et al*., 2018). DDA1 was first identified as a subunit of the plant DDB1-DET1-DDA1 (DDD) complex (Yanagawa *et al*., 2004). DDA1 (Q9FFS4) forms a protein complex with Cul4, DDB1, COP10, and DET1 in *Arabidopsis*, which binds Ub on E2 to the abscisic acid (ABA) receptor protein PYL8 for complete ubiquitination (Irigoyen *et al*., 2014). Studies have also shown that E3 ligase plays a crucial role in plant resistance to disease, including BW (Lee *et al*., 2020; McLellan *et al*., 2020). For instance, both the E3 ligase NtRNF217 in tobacco (Liu *et al*., 2021) and the ATL family gene *StACRE* in potato (Park *et al*., 2012) can positively regulate plant resistance to BW.

Studies have found that many *Solanaceae* crops are not immune to BW (Patil *et al*., 2012). However, eggplant (*Solanum melongena*), as a representative *Solanaceae* crop, is an important vegetable with high BW resistance and sensitivity, making it ideal for BW analysis. Previous research found that eggplant *SmPGH1* is a BW resistance gene (Wang et al., 2020). Besides, eggplant AG91-25 possesses resistance locus *EBWR9*, and the RSSC *ripAX2* gene can induce AG91-25 specific resistance (Morel *et al*., 2018). Xi’ou *et al*. (2015) indicated that the eggplant *RE-bw* gene can interact with the effector Popp2 of RSSC. Qiu *et al*. (2019) also showed that spermidine (SPD) significantly improves eggplant resistance to BW, and SmMYB44 enhances *SmSPDS* expression. However, there is little research on plant resistance to BW at the post-translational level. Moreover, it is unclear whether UPS is involved in the regulation of BW.

Our previous research showed that eggplant NAC transcription factor SmNAC (KM435267) binds to the promoter of SA synthesis gene *ICS1* to inhibit SA accumulation, thereby reducing eggplant BW resistance (Na *et al*., 2016). This study used SmNAC protein as a bait to screen an E3 ubiquitin ligase gene, *SmDDA1b* (GenBank accession number: MZ736671) that interact with SmNAC from the eggplant cDNA library. Besides, this study verified the function of *SmDDA1b* and the relationship between SmDDA1b and SmNAC. Therefore, this research provides new insights into the molecular mechanism by which the SmDDA1b-SmNAC-SA pathway enhances BW resistance.

## Results

### Isolation of eggplant E3 ligase gene *SmDDA1b*

Previous research showed that *SmNAC* can reduce eggplant BW resistance (Na *et al*., 2016). Herein, the N-terminal 417bp base of *SmNAC* containing the NAM domain without self-activation was used as the bait protein to screen the eggplant cDNA library. *SmDDA1b*, 1017bp nucleic acid fragment containing E3 ubiquitin ligase gene was identified. Y2H assay of SmDDA1b and SmNAC was then conducted. AD-*SmDDA1b* and BD-*SmNAC*_*1-417*_ co-transferred to Y2H Gold developed plaque on the four amino acids deficiency medium (SD/-Leu-His-Trp-Ade). The plaque turned blue after the addition of X-α-Gal (Fig. 1A). These results indicate that SmDDA1b can interact with SmNAC.

**Figure 1.**
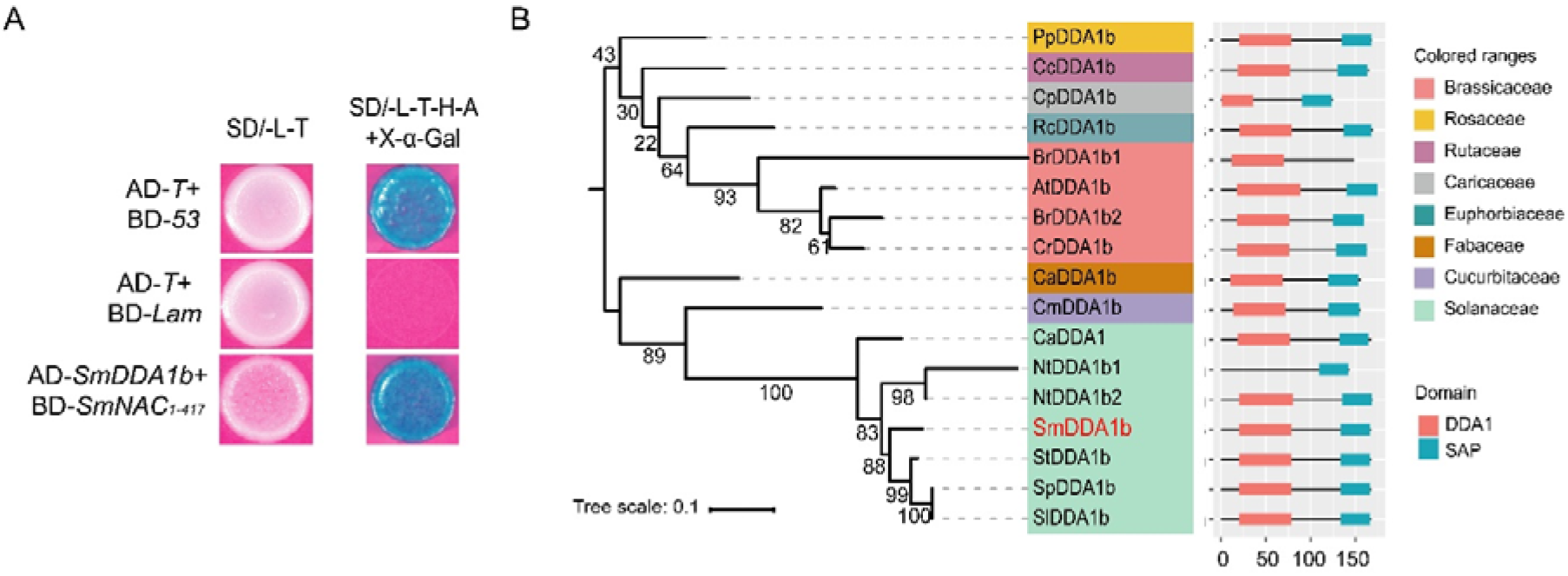
Interaction between SmDDA1b and SmNAC in the yeast two-hybrid (Y2H) system. A, Y2H results of SmNAC and SmDDA1b. The co-transformation of BD-*53*, BD-*Lam* and AD-*T* with Y2H Gold as a positive or negative control. B, SmDDA1b phylogeny analysis results. The number on the branch represents the degree of support, and the maximum value is 100. The genome accession number of the gene is shown in Table S3.

The ORF of *SmDDA1b* (504bp, 167 amino acids, and molecular weight of 18.27 kDa) had DDA1 and SAP domain, belonging to E3 ubiquitin ligase of CRL4 (Supplemental Fig. S1). Fifteen representative dicotyledonous plants with sequenced genomes, including eggplant, were selected for phylogenetic analysis. Each genome was screened to obtain the homologous protein of SmDDA1b (Fig. 1B). They all contained DDA1 and SAP domains except BrDDA1b and NtDDA1b1, indicating that these domains are relatively conserved in dicotyledonous plants and may play a crucial role in the survival and evolution of plants.

### SmDDA1b interacts with SmNAC in the nucleus

Subcellular localization and BiFC experiments were used to determine the position of interaction between SmDDA1b and SmNAC at the subcellular level. The fluorescence microscope showed that the nucleus of tobacco emitted green fluorescence, indicating that *SmDDA1b* is expressed in the nucleus (Supplemental Fig. S4). In the BiFC assay, YNE-*SmDDA1b* and YCE-*SmNAC* produced a small amount of yellow fluorescence in the nucleus when the proteasome inhibitor MG132 was not injected. However, after MG132 injection, the amount of yellow fluorescence in the nucleus increased, indicating that SmDDA1b can interact with SmNAC in the nucleus (Fig. 2A). Therefore, SmNAC could be the ubiquitination substrate of SmDDA1b.

**Figure 2.**
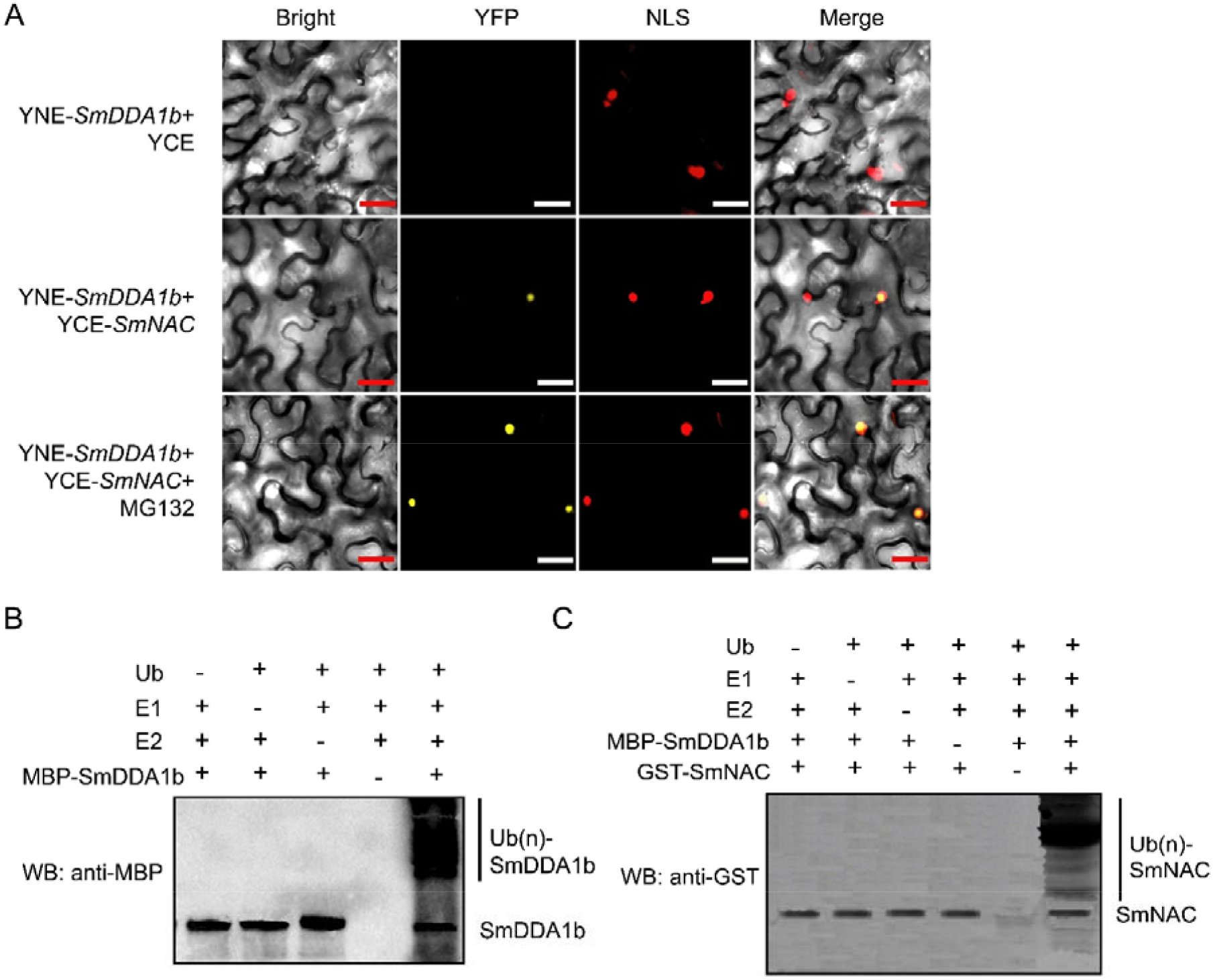
Interaction between SmDDA1b and SmNAC in the nucleus and *in vitro*. A, Bimolecular fluorescence complementation (BiFC) results of SmDDA1b and SmNAC. The proteasome inhibitor MG132 was injected after co-injection of YNE-*SmDDA1b* and YCE-*SmNAC Agrobacterium tumefaciens* in tobacco. YFP indicates yellow fluorescence caused by the interaction between two proteins. NLS indicates the location of the nucleus. The red and white rulers indicate 1 mm. B, The result of E3 ubiquitin ligase activity of SmDDA1b. The symbols “-” and “+” indicate samples not added and those added in the experiment, respectively. A single band represents the SmDDA1b protein, and a ladder-like smear represents the polyubiquitination of SmDDA1b. C, SmNAC ubiquitination via SmDDA1b *in vitro*. A single band represents the SmNAC protein, and the ladder-like smear represents the polyubiquitination of SmNAC.

### SmDDA1b has E3 activity and interacts with SmNAC *in vitro*

A self-ubiquitination experiment was conducted to verify whether SmDDA1b can recognize and degrade SmNAC through the UPS. Polyubiquitination occurred when E1, E2, Ub and MBP-SmDDA1b were all present (Fig. 2B). In contrast, polyubiquitination did not occur in other groups without essential components, indicating that SmDDA1b has E3 ubiquitin ligase activity.

The ubiquitination experiment was then conducted to determine if SmNAC is the target protein of SmDDA1b *in vitro*. (Fig. 2C). A purified GST-SmNAC protein was added into the reaction system containing the above mixture. ZEN-BIOSCIENCE (Chengdu, China) anti-GST antibody was used for western blot (WB) analysis after the reaction was over to detect whether Ub, MBP-SmDDA1b protein, and GST-SmNAC protein were coupled. The ladder-like smear appeared only when all the necessary components were present, indicating that SmNAC polyubiquitination occurred. Therefore, SmDDA1b can ubiquitinate SmNAC *in vitro*.

### SmDDA1b can degrade SmNAC after ubiquitination *in vivo*

Co-IP experiment was performed to further verify the actual ubiquitination of SmDDA1b and SmNAC protein *in vivo*. SmDDA1b-HA and SmNAC-GFP fusion proteins were detected in the protein complex immunoprecipitated with an anti-HA antibody. An indication that SmDDA1b-HA can immunoprecipitate SmNAC-GFP (Fig. 3A). Therefore, SmDDA1b and SmNAC can interact *in vivo*.

**Figure 3.**
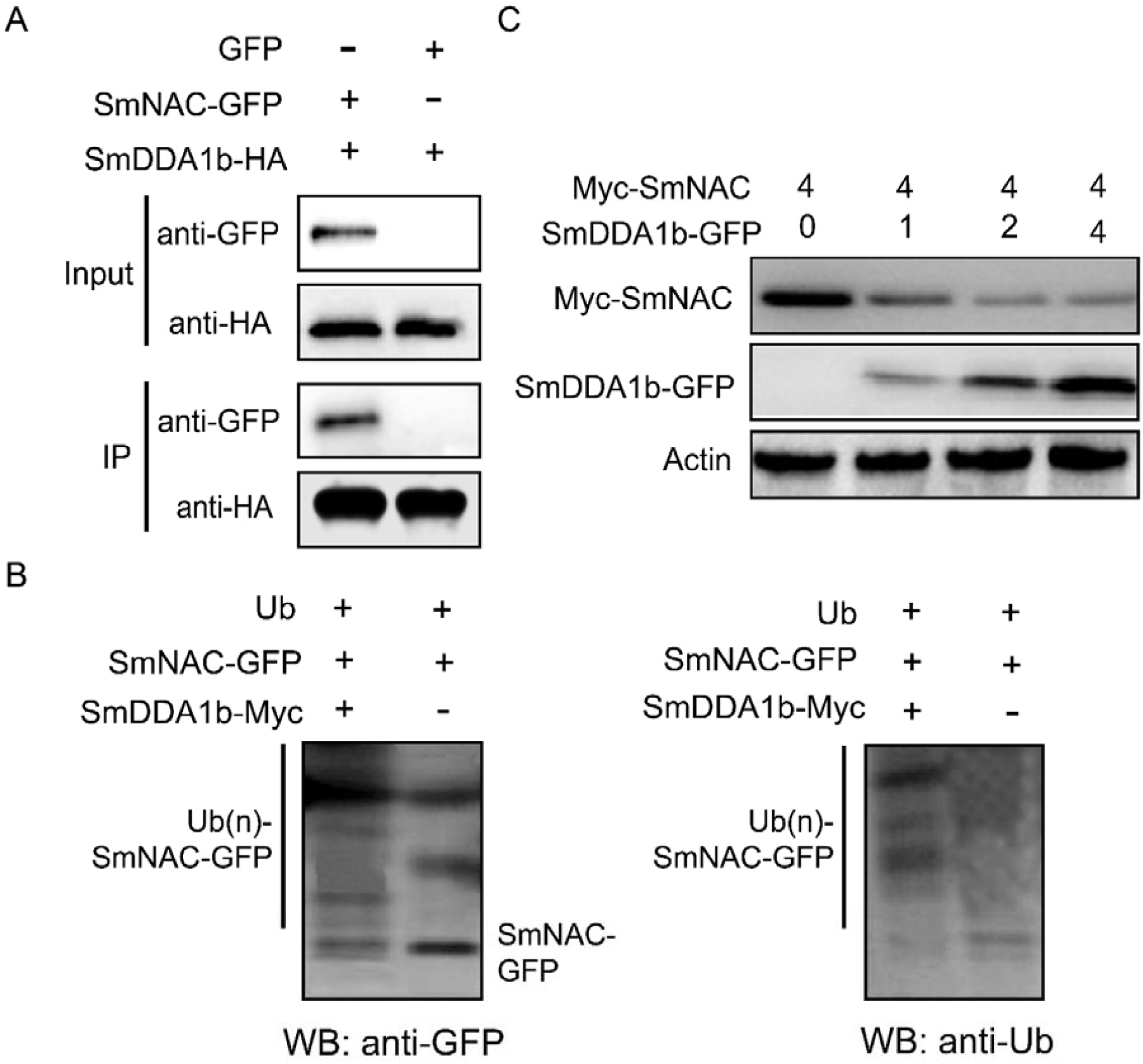
*In vivo* ubiquitination experiment results of SmDDA1b and SmNAC. A, Co-immunoprecipitation (Co-IP) experiment results. Anti-GFP and anti-HA were used for WB detection. B, *In vivo* SmNAC ubiquitination via SmDDA1b. The symbols “-” and “+” indicate samples not added and those added in the experiment, respectively. C, Effect of SmDDA1b-GFP *Agrobacterium tumefaciens* on the expression level of SmNAC protein. Different numbers represent different injection ratios. Anti-Myc and anti-GFP were used for WB detection, and actin was used as a control.

SmDDA1b-Myc and SmNAC-GFP *Agrobacterium tumefaciens* were subjected to tobacco transient expression experiments. Anti-GFP and anti-Ub were used for WB detection. The polyubiquitination band of SmNAC appeared when the two *Agrobacterium tumefaciens* were co-injected. However, anti-GFP showed SmNAC-GFP band, while anti-UB showed no band when only SmNAC-GFP was injected. Therefore, SmDDA1b can modify SmNAC via polyubiquitination *in vivo* (Fig. 3B).

The *Agrobacterium tumefaciens* containing Myc-SmNAC and SmDDA1b-GFP constructs were infiltrated into tobacco leaves for transient expression. The expression of SmNAC protein gradually decreased as the injection ratio of SmDDA1b-GFP protein increased (Fig. 3C). The results further indicate that SmDDA1b can ubiquitinate SmNAC in plants and degrade SmNAC via UPS.

### Expression pattern analysis of *SmDDA1b* in eggplant

Analysis of *SmDDA1b* cDNA sequence was not significantly different between the resistant line E31 and susceptible line E32 (Supplemental Fig. S2). The qRT-PCR assay result showed the expression of *SmDDA1b* was expressed in the roots, stems, and leaves in both of E31 and E32. Notably, the expression level of *SmDDA1b* was higher in E31(R) than in E32(S) (Fig. 4A). After inoculated RSSC into E31 (R) and E32 (S), the qRT-PCR results showed that the expression level of *SmDDA1b* decreased within one hour in both lines. However, *SmDDA1b* expression rapidly increased in E31 (R) after 12 h of inoculation and was not altered in E32 (S) (Fig. 4B). Therefore, the expression level of *SmDDA1b* could be induced by RSSC in the resistant line E31.

**Figure 4.**
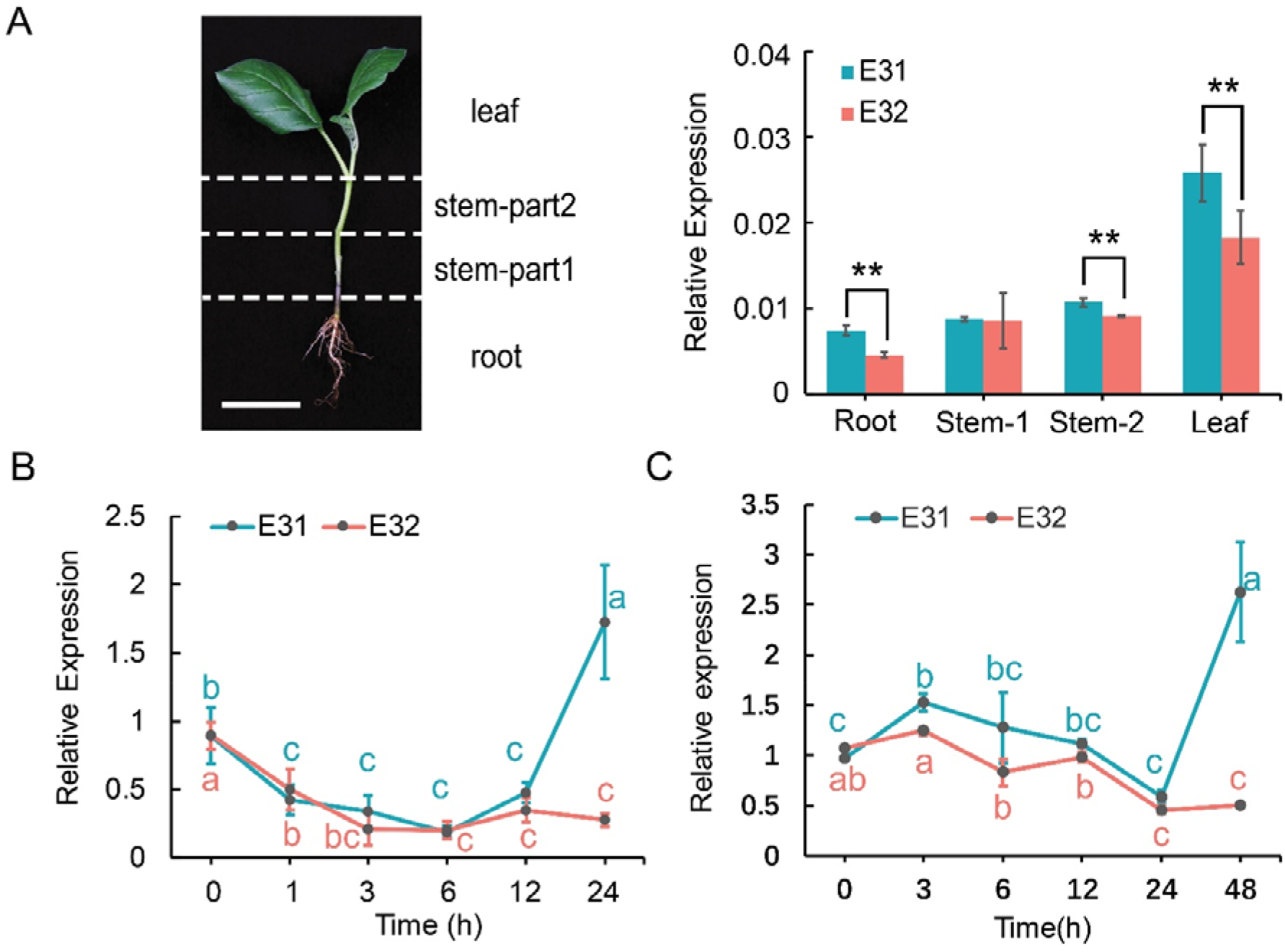
Expression analysis of *SmDDA1b*. A, The relative expression of SmDDA1b in E31 and E32 tissues. The left part shows a schematic diagram of the tissue parts of eggplant seedlings (leaves, upper and lower parts of the stems, and roots). The bar graph shows the relative expression of *SmDDA1b* in the roots, stems-part1, stems-part2, and leaves of eggplant E31 and E32. Data are expressed as mean ± SD (n=3) (*, *p* < 0.05; **, *p* < 0.01, according to Student’s t-test). The white ruler indicates 5 cm. B, The analysis of the relative expression of *SmDDA1b* after 24 h of E31 and E32 inoculation with pathogen. The samples were obtained at 0 h, 1 h, 3 h, 6 h, 12 h and 24 h after treatment. C, The relative expression of *SmDDA1b* after 48 h of E31 and E32 treatment with SA. The samples were obtained at 0 h, 3 h, 6 h, 12 h, 24h and 48 h after treatment. Data are expressed as mean ± SEM of three biological replicates. The letter notation indicates the results of multiple comparisons between the data (*p*<0.05).

At the same time, the expression of *SmDDA1b* in E31 also could be enhance from 0 h to 3 h and rapidly increased after 24 h by SA treatment. However, the expression of *SmDDA1b* in E32 decreased continuously after SA treatment (Fig. 4C). This result indicates that the expression of *SmDDA1b* in E31 could be induced by exogenous SA.

### *SmDDA1b*-silenced eggplant has a decreased BW resistance

VIGS experiment was conducted to verify whether *SmDDA1b* is related to BW resistance. *SmNAC* expression was significantly increased in E31 when *SmDDA1b* expression was reduced (Fig. 5A). Moreover, after inoculation with RSSC, *SmDDA1b*-silenced lines showed clear wilt symptoms and the disease index and morbidity of the SmDDA1b-silenced lines were 70 and 100%, respectively, which were much higher than in control (Fig. 5, B-C). These results indicate that *SmDDA1b* positively regulates eggplant resistance to BW.

**Figure 5.**
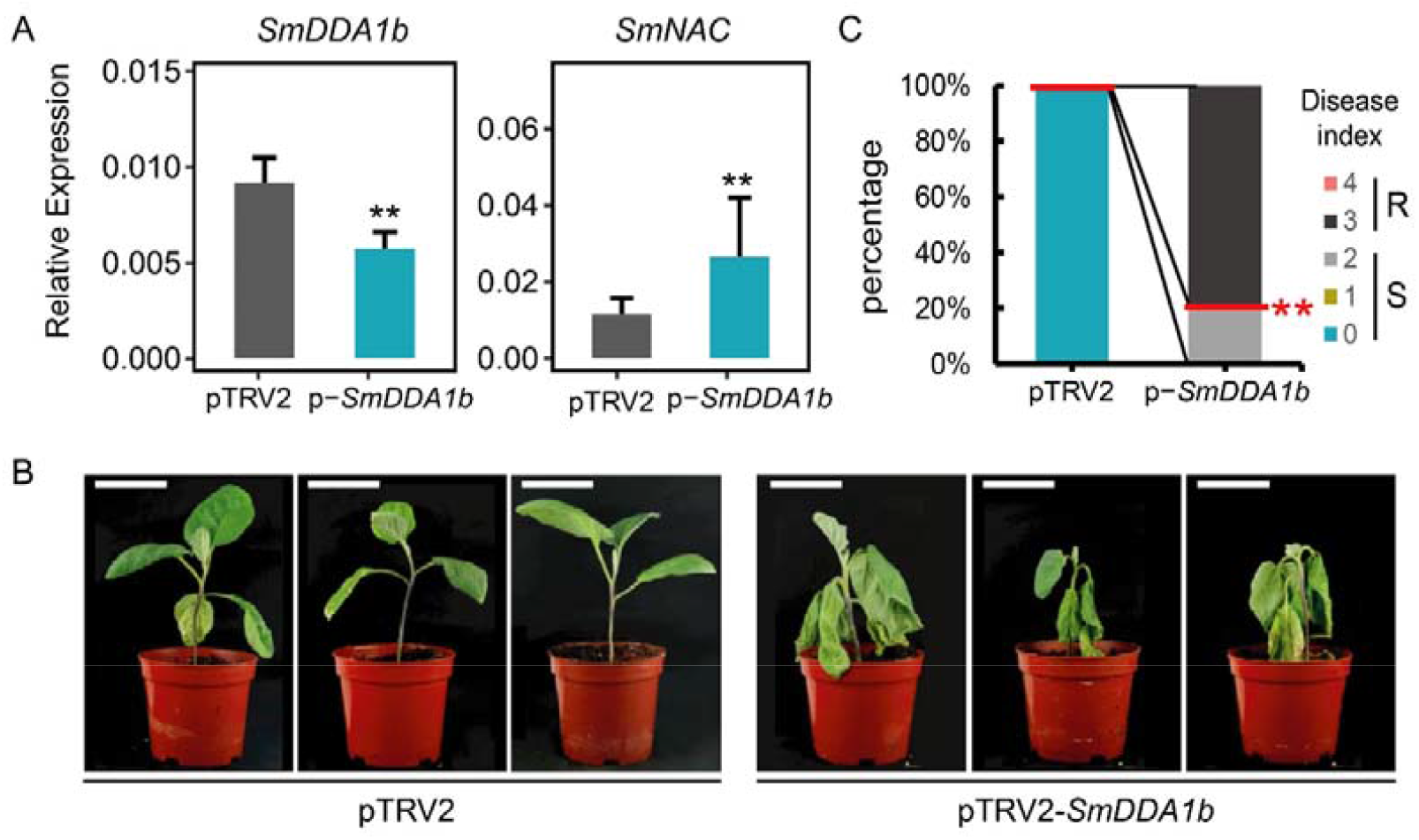
Virus-induced gene silencing (VIGS) experiment results of *SmDDA1b* in E31. A, The relative expression results of *SmDDA1b* and *SmNAC* in *SmDDA1b*-silenced plants. pTRV2 represents the control group, and p-*SmDDA1b* indicates VIGS-treated plants pTRV2-*SmDDA1b*. Data are expressed as mean ± SEM of three biological replicates (**, *p* < 0.01, according to Student’s t-test). B, The phenotype of E31 VIGS control and treated plants infected with *Ralstonia solanacearum* species complex (RSSC) for four weeks. The white ruler indicates 5 cm. C, The results of VIGS and control plant disease index four weeks after inoculation with RSSC. The evaluation scale was as follows: 0= healthy, 1= one or two leaves, wilted, 2= three or more leaves wilted, 3= all leaves wilted, and 4= dead (Qiu *et al*., 2019). The ordinate represents the percentage of the number of plants in each disease level. Ten E31 seedlings were silenced.

### *SmDDA1b* overexpression enhances BW resistance and increases SA content

*SmDDA1b* in tomato cultivar Money Marker was over-expressed to further verify its function. The marker gene *bar* and qRT-PCR were used to obtain 12 T_0_ transgenic tomato seedlings from 60 tissue culture seedlings (Fig. 6A). Three individual plants (OET_0-12_, OET_0-17_, OET_0-31-2_) with good over-expression effects were selected for T_1_ generation propagation (Fig. 6B). Finally, 115 of the 150 individual plants containing *bar* gene were identified using the *bar* marker, and used for subsequent experiments (Supplemental Table S5). After inoculated with RSSC, the WT seedlings were wilted, while the over-expressed seedlings were partially wilted after 7 days and 14 days of inoculation, indicating that over-expressed seedlings are more resistant to BW than the WT plants (Fig. 6, C-D). Besides, the WT and over-expressed seedlings had the same onset time, and both began to show wilting symptoms on the sixth day. However, the morbidity and disease index of WT plants were significantly higher than those of the over-expressed plants after 14 days of inoculation (Fig. 6, E-F; Supplemental Table S6). Taken together, these results indicate that *SmDDA1b* over-expression can increase plant BW resistance.

**Figure 6.**
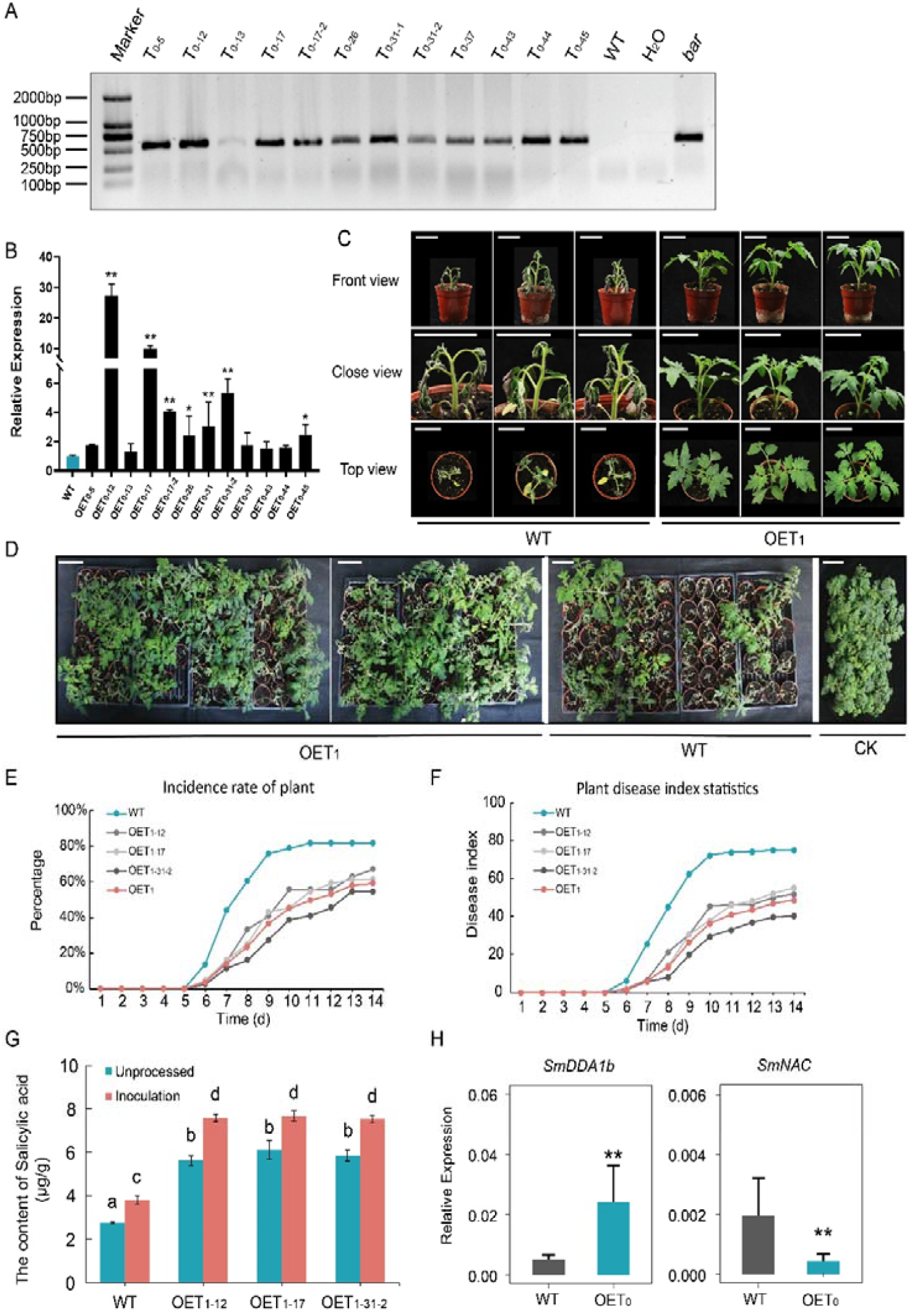
Results of overexpression experiment. A, Detection of the marker gene *bar* on tomato tissue culture seedlings. B, The relative expression of *SmDDA1b* in tomato independent lines obtained from tissue culture. Data are expressed as mean ± SD (n=3) (*, *p* < 0.05; **, *p* < 0.01, according to Student’s t-test). C, The phenotype of wild-type (WT) and T_1_ generation overexpressed seedlings (OET_1_), including front view, close view, and top view on the 7^th^ day after inoculation with RSSC. The white ruler indicates 5 cm. D, The phenotype of WT and OET_1_ on the 14^th^ day post-inoculation with RSSC. CK represents WT and transgenic tomato seedlings that were not inoculated with RSSC. The white ruler indicates 1 dm. E-F, The morbidity statistics and disease index of WT and overexpressed seedlings after 14 days of inoculation. OET_1_ represents the average disease index of three overexpressed lines. G, SA content of WT and overexpressed tomato seedlings inoculated or not inoculated with RSSC. The letters indicate significant differences (*p* <0.05). H, The relative expression of *SmNAC* in *SmDDA1b* overexpressed lines. Data are expressed as mean ± SEM of three biological replicates (*, *p* < 0.05; **, *p* < 0.01, Student’s t-test).

Besides, SA content of WT and *SmDDA1b* over-expressing seedlings inoculated (or not) with RSSC was determined. SA content was significantly lower in WT than in over-expressed seedlings before and after inoculation, indicating that over-expression of *SmDDA1b* in tomato can increase SA content. The SA content of both WT and over-expressed lines was significantly increased after inoculation (Fig. 6G). Therefore, *SmDDA1b* can regulate plant resistance to BW by altering SA content. Moreover, the expression of *SmNAC* was significantly decreased in *SmDDA1b* over-expressed lines compared with the WT (Fig. 6H), which was consistent with the study that SmDDA1b could degrade SmNAC through UPS.

### SmDDA1b indirectly and positively regulates *ICS1* and SA pathway gene expression

ICS1 can synthesize SA, in order to detect whether SmDDA1b affects ICS1, the expression of *ICS1* (NM_001247865.1) in *SmDDA1b* over-expressed and VIGS plants was analyzed. In the over-expressed plants, the expression of *ICS1* was increased, and in the VIGS plants, the *ICS1* expression was decreased, indicating that SmDDA1b can positively regulates the expression of *ICS1* (Fig. 7A). Besides, Y2H and BiFC results of SmDDA1b and ICS1 show that SmDDA1b can not directly target ICS1 (Supplemental Fig. S5). The results deduced that *SmDDA1b* degrade *SmNAC* to positively increase activity of *ICS*1.

**Figure 7.**
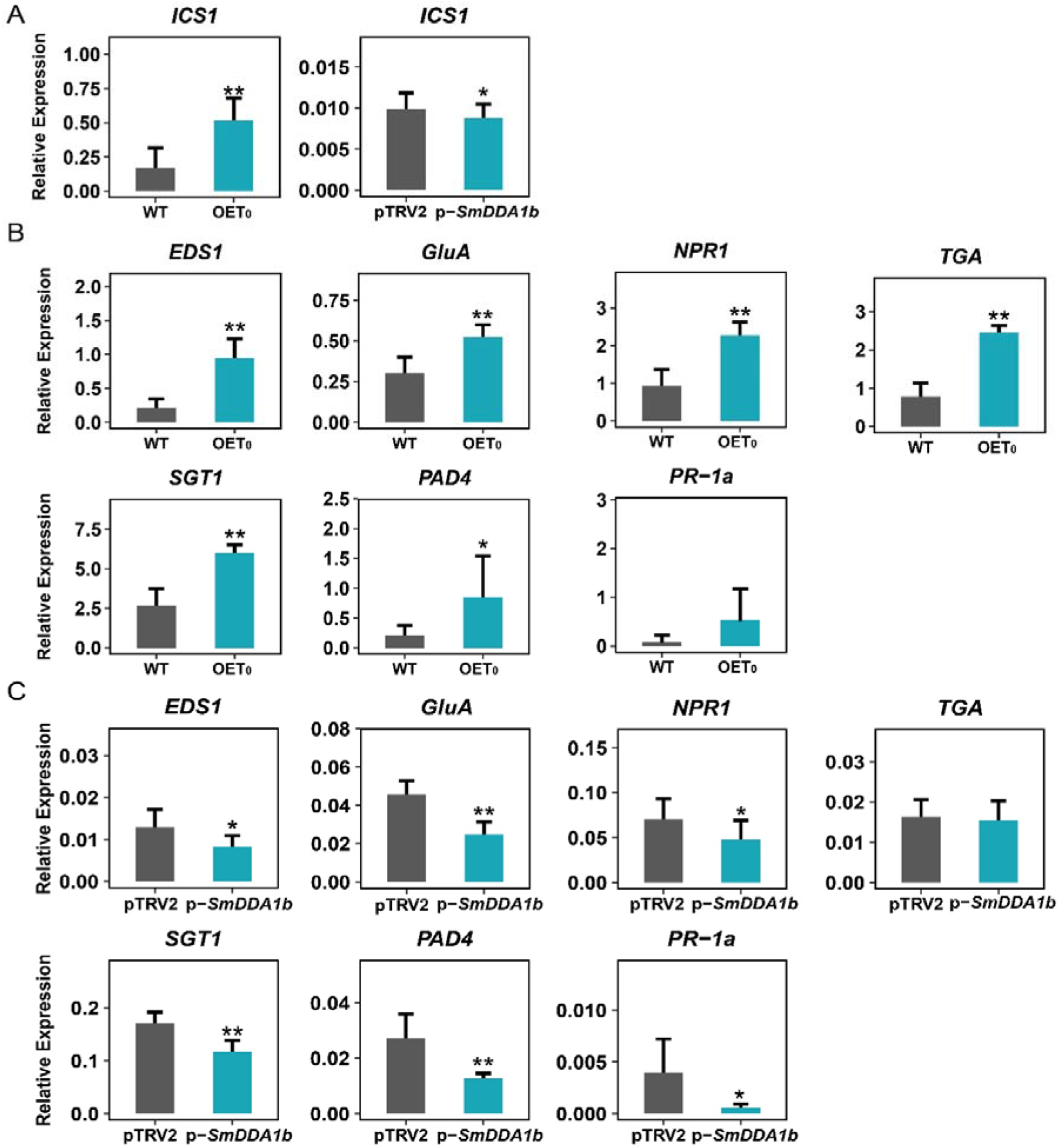
Expression of *ICS1* and salicylic acid (SA) pathway signal-related genes in *SmDDA1b* overexpressed and virus-induced gene silencing (VIGS) plants. A, Expression of *ICS1* in *SmDDA1b* overexpression and VIGS plants. B, Expression of SA pathway signal-related genes in overexpressed plants. OET_0_ represents the T_0_ generation overexpressed plants. and WT represents the wild-type plants. C, Expression of SA pathway signal-related genes in VIGS plants. pTRV2 represents control plants, and p-*SmDDA1b* represents VIGS-treated plants pTRV2-*SmDDA1b*. Data are expressed as mean ± SEM of three biological replicates (*, *p* < 0.05; **, *p* < 0.01, Student’s t-test).

This study also analyzed the expression of hormone signal pathway-related genes in *SmDDA1b* over-expressed and VIGS plants. *EDS1* (AY679160.1), *GluA* (M80604), *NPR1* (NM_001247633.1), *TGA* (GQ386946.1), *SGT1* (NM_001247758.1), *PAD4* (AY753546.1) and *PR-1a* (M69247) were selected to assess if SA pathway signal-related genes can regulate BW resistance. The expression of SA pathway signal-related genes was significantly increased in the over-expressed plants except for *PR-1a* (compared with the control) (Fig. 7B). In contrast, the expression of SA pathway signal-related genes was significantly decreased in VIGS plants except for TGA (compared with the control) (Fig. 7C), indicating that *SmDDA1b* positively regulates the expression of SA pathway signal-related genes.

### SmNAC binds to *SmDDA1b* promoter and significantly represses the promoter activity

In previous studies, *SmNAC* over-expression lines have shown decreased expression of *SmDDA1b* (Fig. 8A). This suggests that the activity of *SmDDA1b* promotor might be directly down-regulated by SmNAC. In order to verify this hypothesis, the *SmDDA1b* promoter sequence was obtained from eggplant genome (Barchi *et al*., 2021), and the elements of the promoter were predicted by PlantPAN 3.0 (Supplemental Fig. S6). It was predicted that the *SmDDA1b* promoter contained 24 NAC element binding sites and these sites are mostly distributed in the region of - 500 to - 1500, indicating that SmNAC may bind to *SmDDA1b* promoter, then the promoter was isolated and cloned (Fig. 8B; Supplemental Table S7).

**Figure 8.**
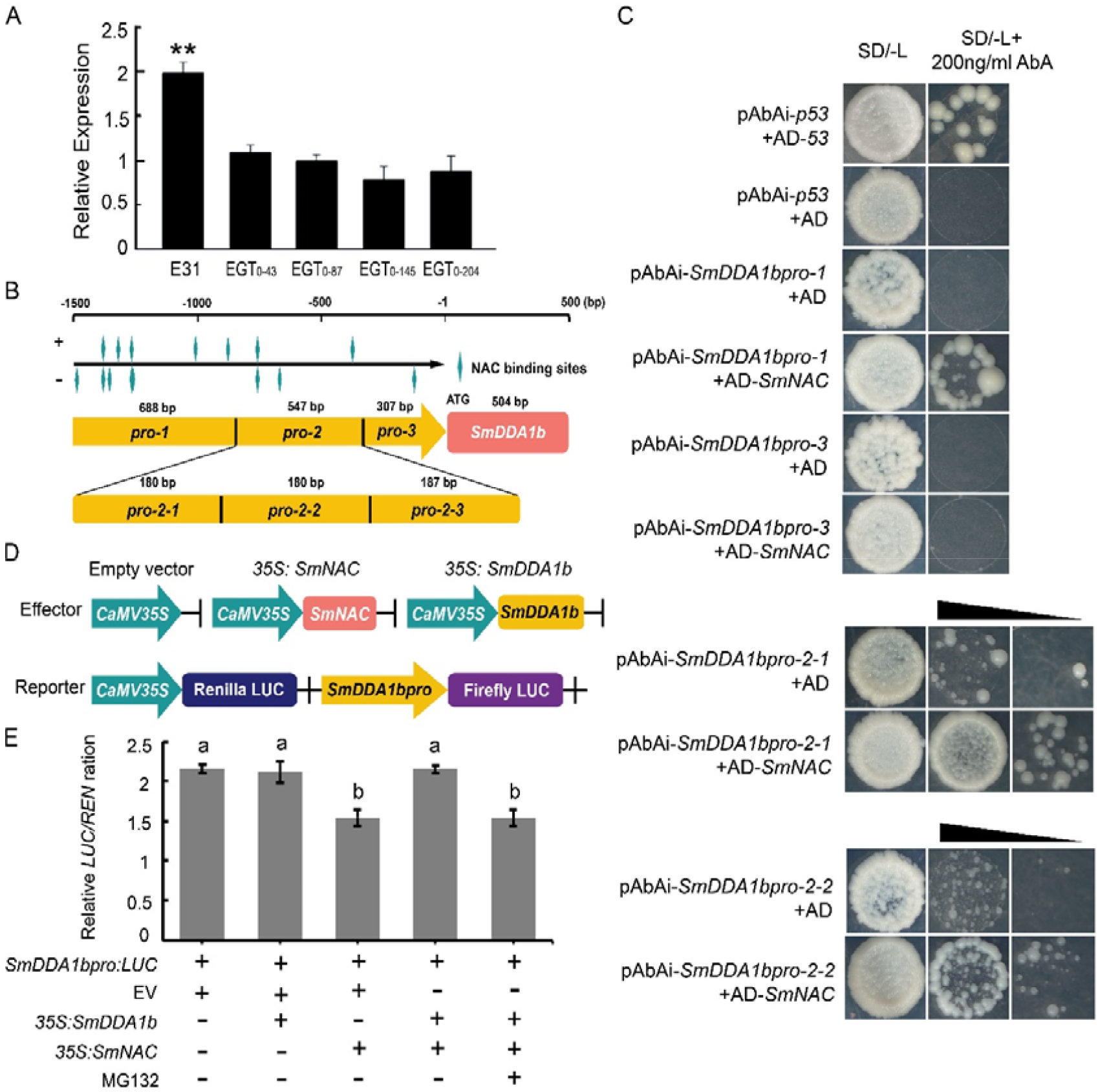
SmNAC binds to the *SmDDA1b* promoter and represses the expression of *SmDDA1b* gene. A, The relative expression of *SmDDA1b* in *SmNAC* over-expressed lines. E31 indicates wild-type, and EGT_0-43_, EGT_0-87_, EGT_0-145_, EGT_0-204_ represent T_0_ generation over-expressed plants. Data are expressed as mean ± SEM of three biological replicates (**, p < 0.01, Student’s t-test). B, Schematic diagram of *SmDDA1b* gene, promoter and NAC element binding sites. “+” represents the sense strand and “-” represents the antisense strand. The promoter of *SmDDA1b* was divided into three segments for Y1H assay. The second segment had self-activation and it was divided into three segments for Y1H with SmNAC transcription factor. *SmDDA1bpro-1*: -855 to-1542 bp, *SmDDA1bpro-2*: -308 to -854 bp (*SmDDA1bpro-2-1*: -675 to -854 bp, *SmDDA1bpro-2-2*: -495 to -674 bp, *SmDDA1bpro-2-3*: -308 to -494 bp), *SmDDA1bpro-3*: -1 to -307 bp. C, SmNAC binds directly to the SmDDA1b promoter. The co-transformation of AD-*53*, AD and pAbAi-*p53* with Y1H Gold as a positive or negative control. The triangle indicates the yeast concentration from high to low. D, A schematic representation of the double reporter and effector plasmid used. The double-reporter plasmid contained *SmDDA1b* promoter fused with the sequence encoding LUC luciferase and REN luciferase driven by *CaMV35S* promoter. The effector plasmid contained *SmNAC* driven by a *CaMV35S* promoter. E, SmDDA1b attenuates the inhibitory effect of SmNAC on SmDDA1b promoter. The regulation of transcription factor on promoter activity is based on the ratio of LUC to REN. The symbols “-” and “+” indicate samples not added and those added in the experiment, respectively. EV represent empty vector. Data are expressed as mean ± SEM of three biological replicates. The letter notation indicates the results of multiple comparisons between the data (*p*<0.05).

Due to the self-activation of *SmDDA1b* promoter, the promoter was divided into three segments for yeast one-hybrid (Y1H) assay, named *SmDDA1bpro-1, SmDDA1bpro-2* and *SmDDA1bpro-3*, respectively, and divided *SmDDA1bpro-2* into *pro2-1, pro2-2, pro2-3* (Fig. 8B), among them, *SmDDA1b pro2-3* has self-activation (Supplemental Fig. S7). The results of Y1H showed that there was no significant difference between pAbAi-*SmDDA1bpro-3* and AD-*SmNAC* co-transformed to Y1H Gold and the control, no yeast plaque grew on the Leucine deficiency medium (SD/-L) added with 200ng/ml Aureobasidin A (AbA). However, the pAbAi-*SmDDA1bpro-1/ pro2-1*/*pro2-2* and AD-*SmNAC* co-transformed to Y1H had significant yeast plaque growth on SD/-L added with 200ng/ml ABA compared with the control, and the results remained unchanged after dilution (Fig. 8C). The results showed that SmNAC could bind to *SmDDA1bpro-1, pro-2-1* and *pro-2-2* regions.

In order to further verify the regulatory effect of SmNAC on the activity of *SmDDA1b* promoter, we carried out dual-luciferase assay. We found that the ratio of LUC / REN in the treatment group was significantly lower than that in the control group, indicating that SmNAC represses the transcription of *SmDDA1b* (Fig. 8, D-E). In addition, after *Agrobacterium tumefaciens 35S: SmDDA1b, 35S: SmNAC* and *SmDDA1bpro: LUC* were co-injected into tobacco, the ratio of LUC / REN increased and the inhibitory effect of SmNAC on the promoter was eliminated. After MG132 injection, the ratio of LUC/ REN decreased and the inhibitory effect of SmNAC on promoter was restored (Fig. 8E). The results showed that SmDDA1b could degrade SmNAC through UPS, and the increase of SmDDA1b content could weaken or even remove the inhibitory effect of SmNAC on *SmDDA1b* promoter.

## Discussion

E3 ubiquitin ligase and NAC (NAM-ATAF-CUC1/2) transcription factors are crucial in plant disease resistance. The E3 ligase has a complex plant disease resistance regulation, including positive and negative regulation. Besides, some interact with pathogens or post-translational modifications of other proteins to directly or indirectly regulate plant disease resistance (Miao et al., 2016; Wang et al., 2020; Karki et al., 2021). For instance, MIEL1 is a RING-type E3 ligase, negatively regulating defense response in *Arabidopsis* (Marino *et al*., 2013). The E3 ligase NbUbE3R1 positively regulates the immune response in tobacco. Furthermore, the replicase of *Bamboo mosaic virus* (BaMV) could be a substrate of NbUbE3R1 (Chen *et al*., 2019). RING-type E3 ligase, VIM5, can target and degrade DNA methyltransferases MET1 and CMT3 through the 26S proteasome. *Beet severe curly top virus* can induce *VIM5* expression and activate the *C2* and *C3* genes of the geminivirus to make the plant susceptible (Chen et al., 2020).

NAC, as a unique family of transcription factors in plants, is essential at multiple levels of transcription, post-transcriptional and post-translational modification (Zhu et al., 2016; Zhang et al., 2018; Li et al., 2019; Liu et al., 2020). Studies have shown that E3 ligase can interact with NAC transcription factors (Yoshii *et al*., 2010). SINA protein also has E3 ubiquitin ligase activity (Wang *et al*., 2018). The RING-type E3 ligase SINAT5 can ubiquitinate the NAC transcription factor AtNAC in *Arabidopsis* (Xie *et al*., 2002). Miao *et al*. (2016) indicated that SINA can recognize and degrade NAC1 in tomato through the UPS, negatively regulating the role of plant defense signals.

Herein, SmDDA1b ubiquitinated SmNAC *in vivo* and *in vitro*, promoting its degradation through UPS. However, E3 ubiquitin ligase and NAC transcription factor are only one aspect of this mechanism. Besides targeting SmNAC, SmDDA1b may also target other factors that can negatively regulate eggplant BW resistance via ubiquitination and degradation. Therefore, future research should focus on the regulation network of resistance to BW.

Plants balance their gene expression and control the role of E3 ligase and NAC transcription factors when there is no biological stress. Previous studies have shown that transcription factor, the target protein of E3 ligase, can also bind to the promoter element of E3 ligase to control the expression activity of E3. Tong *et al*. (2021) found that *Populus* U-box E3 ligase PalPUB79 degraded PalWRKY77 through ubiquitination, at the same time, PalWRKY77 can bind to the PalPUB79 promoter to represses the expression of PalPUB79 under normal conditions. In tartary buckwheat (*Fagopyrum tataricum*, TB), the E3 ligase FtBPM3 target protein FtMYB11 can also bind to the *FtBPM3* promoter and directly represses the expression of *FtBPM3* gene (Ding *et al*., 2021). Herein, *SmNAC* bind to the promoter element of *SmDDA1b* and negatively regulate *SmDDA1b*. This regulation effect can inhibit the degradation of *SmNAC* and thus maintaining the stability of SmNAC protein and E3 ligase.

SA and SA signaling pathway genes regulate each other. EDS1 can cause the initial accumulation of SA and interact with PAD4 to cause further accumulation of SA, which is located upstream of the signaling pathway (Feys et al., 2001; Wildermuth et al., 2001; Cui et al., 2017). NPR1 acts downstream of the SA signaling pathway and directly affects the SA content (Ding *et al*., 2018). SA also promotes the expression of SA signal pathway genes through positive feedback, thereby rapidly amplify SA signals (Wiermer et al., 2005; Wu et al., 2012; Oh et al., 2014). Herein, *SmDDA1b* overexpressed caused the increase of SA content and the relative expression level of signal genes in SA pathway, while *SmDDA1b* was silenced, the expression of signal genes was decreased, indicating that our research results are consistent with previous research results.

Different signaling pathways interact to form complex signal networks. Plants regulate different defense signal transduction pathways through this signal network to obtain higher stress tolerance (Derksen et al., 2013; Checker et al., 2018). Recent studies have also shown that E3 ligase CUL3^BPM^ can target MYC2, MYC3, and MYC4, reduce the abundance of MYC protein, and regulate the JA pathway (Chico *et al*., 2020). Moreover, RING-type E3 ligase KEG can positively regulate the expression of the JA pathway signal-related gene *JAZ12* (Pauwels *et al*., 2015). SA and JA signals are mutually antagonistic (Adams and Spoel, 2018; Nakano and Mukaihara, 2018). Other hormones may also regulate *SmDDA1b* and should be further verified.

Herein, *SmDDA1b* was first decreased, then increased in the resistant plants after inoculation with RSSC. *SmDDA1b* first decreased, then leveled off in the susceptible plants after inoculation with RSSC. The inhibition may be related to the immune response of plants and pathogenic effectors. The innate immune system of plants (the immune response stimulated by pathogen-related molecular patterns, pattern-triggered immunity (PTI), and effector proteins, effector-triggered immunity (Hernández and Sanan-Mishra)) respond during pathogen invasion. PTI is a nonspecific basic defense response, while ETI is a specific response induced by the plant resistance protein to recognize pathogens (Nakano *et al*., 2017). During pathogen invasion, plants first induce PTI, after which the pathogen releases effectors to inhibit PTI, decreasing *SmDDA1b* expression in both resistant and susceptible plants. Plants then exert an ETI response to inhibit effectors, increasing the *SmDDA1b* expression in disease-resistant plants.

E3 ubiquitin ligase may target the pathogenic effector. Studies have shown that UPS can specifically recognize pathogenic effectors in plants and play a role in plant-pathogen interactions (Zhang et al., 2011; Li et al., 2014; Zhang et al., 2020). Drugeon and Jupin (2002) showed that UPS can target the motor protein 69k of *turnip yellow mosaic viru*s (TYMV) and regulate its activity *in vitro*. The RING-type E3 ligase NtRFP1 can mediate the degradation of geminivirus-encoded βC1 in tobacco (Shen *et al*., 2016). RSSC contains various secretion systems but mainly exerts its effects through the type III secretion system (T3SS). T3SS can influence the host to cause plant diseases or hypersensitivity response (HR) (Lindgren, 1997; Poueymiro and Genin, 2009). Therefore, E3 can target the virulence genes and effectors of RSSC and degrade them via ubiquitination to improve eggplant resistance, based on the specificity of SmDDA1b for the defense response of RSSC. However, further studies are needed to verify the above phenomenon.

## Conclusions

In summary, this study constructed a SmDDA1b-SmNAC-SA pathway regulatory module and showed that SmDDA1b can degrade SmNAC through UPS to enhance BW resistance. Under normal conditions, SmNAC represses the transcription of both *SmDDA1b* and *ICS1* to maintain the immune balance of plants, endogenous SA levels are low in eggplant (Fig. 9A). However, *SmDDA1b* gene was up-regulated after inoculating disease-resistant plants with RSSC, thus decreasing SmNAC expression and the inhibitory effect on *SmDDA1b* decreased, and then increasing *ICS1* expression, SA content and BW resistance (Fig. 9B). Besides, SmDDA1b could not target ICS1 directly.

**Figure 9.**
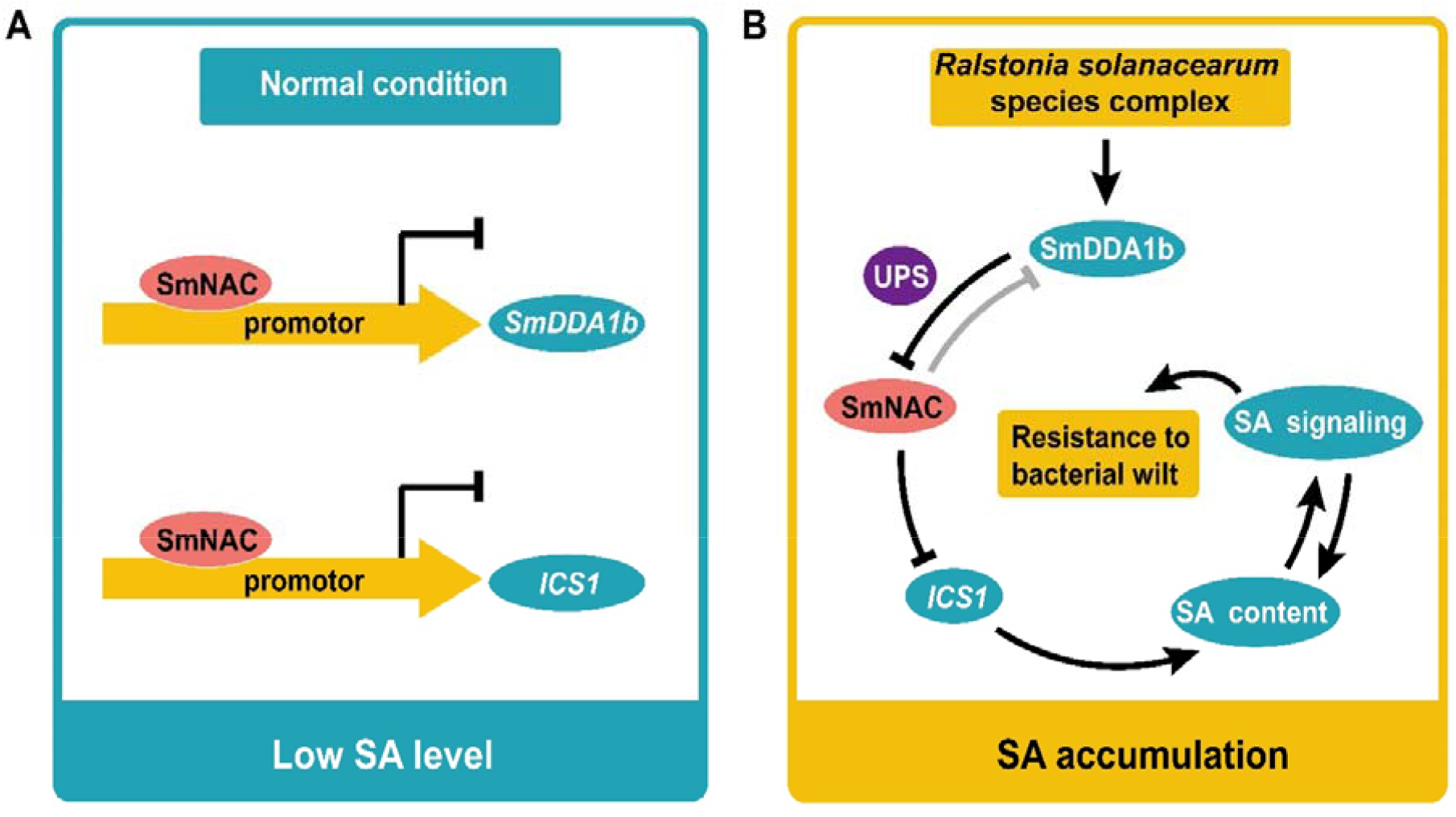
A proposed model for the action mechanism of SmDDA1b. A, SmNAC inhibited the expression level of *SmDDA1b* and *ICS1* under normal condition. B, The regulation module of SmDDA1b improving plant resistance to BW. The expression of SmDDA1b increased under RSSC stress weakens the SmNAC-mediated inhibition of *SmDDA1b*. Moreover, SmNAC-mediated inhibition of *ICS1* was relieved by SmDDA1b-mediated ubiquitination of SmNAC, and thus SA accumulation.

Plant defense against pathogens involves complex mechanisms and many aspects. At the same time, the importance of SA for plant disease resistance is self-evident. Therefore, future researches should explore: screening of E3 ubiquitin ligase genes that can interact with SmNAC except *SmDDA1b*; whether SmDDA1b can interact with pathogenic effector proteins to degrade it via ubiquitination; SmDDA1b can ultimately regulate the content of SA, whether SmDDA1b has the function of resisting other diseases, or regulating plant resistance and growth and development.

## Materials and Methods

### Experimental materials

This study used two eggplant inbred lines, E31 (resistant to BW, R) and E32 (susceptible to BW, S) (Supplemental Fig. S3; Supplemental Table S4). Tomato, tobacco, and RSSC strain used included Money Maker, *Nicotiana benthamiana*, and GMI1000, respectively.

### Data analysis

Total RNA isolation, complementary DNA (cDNA) synthesis, and real-time reverse transcription-PCR (qRT-PCR) were performed using previously described methods (Qiu *et al*., 2019). The relative expression amount was calculated using the 2^-Δct^ and 2^-ΔΔct^ methods (Livak and Schmittgen, 2001). *18SrRNA* was used as the reference gene. qRT-PCR primers are listed in Table S2.

### Phylogenetic analysis and sequence alignment

DDA1 containing sequences of 15 dicotyledonous plants (including eggplant) were obtained by scanning whole-genome protein sequences (Supplemental Table S3) from the NCBI RefSeq database (O’Leary *et al*., 2016) using Hmmserch v3.3 (Eddy, 1998). Mafft v7.455 was used to align sequences (Katoh and Standley, 2013). Iqtree v1.6.12 (Nguyen *et al*., 2015) was then used to construct a phylogenetic tree. Sequence alignment was performed in DNAMAN (version 7.0; Lynnon Biosoft, Quebec, Canada).

### Subcellular localization analysis

The full-length coding sequence of *SmDDA1b* without the stop codon was cloned into the *Age* I site of the pEAQ-EGFP vector (Sun *et al*., 2020). The recombinant vector was introduced into *Agrobacterium tumefaciens* (strain GV3101(pSoup)) and mixed with *Agrobacterium tumefaciens* with DsRed protein (v: v, 1: 1) (Sun et al., 2020). The nuclear-localized signal (NLS) was fused to DsRed as a nuclear marker. The mixture was injected into *Nicotiana benthamiana*, then incubated at 22 °C for three days in the dark. A confocal fluorescence microscope (Carl Zeiss, Germany) was used to visualize green fluorescent protein (GFP) fluorescence. The experiment was repeated at least thrice. The primers are listed in Table S1.

### Pathogen inoculation

Inoculation with RSSC was performed as described in our previous study with some modifications (Qiu *et al*., 2019). Briefly, RSSC was grown in a TTC medium (Lemessa and Zeller, 2007) at 30 °C for two days. The concentration of the inoculum was then determined using a spectrophotometer, and OD_600_ was adjusted to 0.6. Four- to five-day-old seedlings were inoculated by wounding the roots, then incubated in the bacterial suspension for 20 min before transplanting. The entire experiment was conducted under control conditions (30 °C, 16 h of light, and 24 °C, 8 h of dark). The control group was treated with water. Samples (three biological replicates each) were taken at 0 h, 1 h, 3 h, 6 h, 12 h and 24 h.

### Hormone treatment

The four- or five- euphyllas- old eggplant seedlings were treated with 1mM SA and sprayed every 12 hours (Jia et al., 2013; Hussain et al., 2018; Mahesh and Sharada, 2018). The control group was treated with water, then planted at 26 °C, 16 h light, and 22 °C, 8 h dark. Samples (three biological replicates each) were obtained at 0 h, 2 h, 6 h, 12 h, 24h and 48 h.

### Virus-induced gene silencing (VIGS) assays

The specific fragments of about 300 bp from *SmDDA1b* were cloned into *Eco*R I and *Sma* I sites of the pTRV2 vector. pTRV2-*SmDDA1b*, pTRV2, and pTRV1 vectors were then transferred into the *Agrobacterium tumefaciens* strain GV3101. pTRV1 mixed with pTRV2 or pTRV2-SmDDA1b (v: v, 1: 1) were infiltrated into four- or five-day-old seedlings using a 1 mL needleless syringe. After injection, the samples were treated at 16 °C in the dark for one day and then planted normally for 1-2 weeks (26 °C, 16 h light, 22 °C, 8 h darkness). Each treatment had at least 10 biological replicates. Primers are shown in Table S1.

### Construction of the *SmDDA1b* overexpression vector and transformation procedures

The forward primer 5’-gagaacacgggggactctagaATGGAGGATACCTCATCATCCATT-3’ and the reverse primer 5’-gtggctagcgttaacactagtTCATGTGTCCCCCCTTAACCG-3’ were used to amplify the full-length *SmDDA1b* and cloned into *Xba* I and *Spe* I of the pCAMBIA-1380 vector, then transfected into the *Agrobacterium* strain GV3101. The resulting overexpression vector, pCAMBIA-1380-*SmDDA1b*, containing the CaMV35S promoter, Nos terminator, and the *bar* marker gene (5’-end primer ATGAGCCCAGAACGACGCCCG, 3’-end primer TTAGATCTCGGTGACGGGCAGGACC) were then transformed into tomato Money Marker. The transgenic plants were generated as described by Qiu *et al*. (2016).

### Salicylic acid (SA) extraction and quantification

The leaves from *SlDDA1b*-overexpressed lines and non-transgenic lines (wild type) before and after inoculation with RSSC were used for SA extraction and determination. SA extraction and quantification were performed as previously described by Ma *et al*. (2018).

### Yeast two-hybrid (Y2H) assay

The full-length *SmDDA1b* was cloned into *Eco*R I and *Bam*H I sites of the pGADT7 vector. The other genes or fragments, including the N-terminal 417 bases of *SmNAC*, which did not exhibit autoactivation, and the full-length ICS1 without the stop codon, were ligated into the pGBKT7 vector to generate baits. The specific primers and corresponding construction vectors are shown in Table S1. The experiment was conducted following the manufacturer’s instructions (Cat. No. 630489; Clontech, Mountain View, CA, USA).

### Bimolecular fluorescence complementation (BiFC) analysis

The full-length *SmDDA1b* without the stop codon was cloned into *Sal* I and *Bam*H I sites of the pSPYNE-35s / pUC-SPYNE (YNE) vector containing the N-terminal of yellow fluorescence protein (YFP). The other genes without the termination codon were ligated into the pSPYCE-35s / pUC-SPYCE (YCE) vector containing the C-terminal of YPF. The construct was introduced into *Agrobacterium tumefaciens* GV3101(pSoup). The samples were mixed with *Agrobacterium tumefaciens* harboring DsRed protein (v: v: v, 1: 1: 1) injected into *Nicotiana benthamiana*, then planted at 22 °C for three days in the dark. Proteasome inhibitor MG132 (50 μM) was injected (Marques *et al*., 2009). A confocal fluorescence microscope (Carl Zeiss, Germany) was used to visualize GFP fluorescence. The experiment was repeated at least thrice. The primers are listed in Table S1.

### *In vitro* ubiquitination

The full-length *SmNAC* and *SmDDA1b* were cloned into pGEX-4T and pMAL-c2X vector at *Sal* I and *Xho* I sites, respectively. The constructs were then transferred to BM Rosetta (DE3). The ubiquitination reaction mixture (30 μL) contained 600 ng GST-SmNAC protein, 600 ng MBP-SmDDA1b protein, 20× prepared reaction buffer (1 mM ZnCl_2_, 200 mM MgCl_2_, 1 M Tris-HCl, 20 mM ATP, 4 mM DTT, 200 mM creatine phosphate), 0.1 unit creatine kinase (Sigma, USA), 50 ng E1 (Boston Biochem, USA), 250 ng E2 (Boston Biochem, USA). Sterile water was added to make up the solution to 30 μL. The reaction was conducted at 37 °C for 60-90 min. A 7 μL of 5×Loading buffer was added to a 95 °C water bath for 5 min. The sample was then centrifuged at 10000 rpm for 1 min to obtain supernatant for SDS-PAGE electrophoresis. Western blot was conducted as described by Na *et al*. (2016). anti-GST (ZEN-BIOSCIENCE, China) antibody was used. Primers are listed in Table S1.

### *In vivo* ubiquitination

The full-length *SmNAC* and *SmDDA1b* were constructed into *Hind* III and *Sal* I sites of pC1307-35S-Myc and pC2300-35S-GFP vectors, respectively. The extracted plasmids were transferred into GV3101. The supernatant of *Agrobacterium tumefaciens* (OD_600_= 0.6) was resuspended in infection buffer (10 mM MgCl_2_, 10 mM MES (pH 5.6), 100 μM AS) solution and allowed to stand for 2-5 h. Myc-SmNAC was then injected into *Nicotiana benthamiana*. SmNAC expression was unchanged (OD_600_=0.4), while the injection ratio of SmDDA1b was gradually increased (OD_600_=0-0.4). The samples were incubated at 22 °C for 2 d, and then western blot analysis was conducted.

The *SmNAC*-GFP and *SmDDA1b*-Myc vectors were constructed and transformed into GV3101. The *Agrobacterium tumefaciens* (OD_600_= 0.6) was resuspended in infection buffer and allowed to stand for 2-4 h. The injection of tobacco with *Agrobacterium tumefaciens* liquid of *SmDDA1b*-Myc and *SmNAC*-GFP was set as control. Only *SmNAC*-GFP was injected in the treatment group. Western blot analysis was conducted after incubation at 22 °C for two days. The antibodies used were anti-GFP and anti-Ub. Primers are shown in Table S1.

### Co-immunoprecipitation (Co-IP) assay

The full-length *SmDDA1b* was cloned into pAC004-HA vector to produce SmDDA1b-HA antibody, and *SmNAC* was cloned into pAC402-GFP vector. The *Agrobacterium tumefaciens* with GFP-tagged empty plasmid, *SmNAC* recombinant plasmid, and HA-tagged *SmDDA1b* recombinant vector was diluted in infection buffer to OD_600_=1.2. pAC402-X (Vec or SmNAC) and pAC004-*SmDDA1b* (v:v, 1: 1) was then added to the sample to co-infect tobacco. The samples were obtained after 48 h of infection, then lysed to obtain input. Western blot was used to detect the expression. GFP-Trap agarose magnetic beads with immobilized GFP antibody were used to incubate the protein at 4 °C for 1 h. The beads were put on a magnetic stand for 1 min and washed twice with the wash buffer (50 mM Tris-HCl, 5 mM EDTA, 250 mM NaCl, 1 mM PMSF, 10% glycerol, pH 7.5). A loading buffer was added, then boiled and centrifuged to obtain supernatant (IP sample). Western blot was used to check. Primers are listed in Table S1.

### Promoter isolation and element prediction

Download the eggplant genome data from Sol Genomics NetWork (https://solgenomics.net) (Barchi *et al*., 2021). TBtools v1.09852 (Chen et al., 2020) was used to blast the genome to find out the *SmDDA1b* gene, the 1474bp fragment before the *SmDDA1b* gene was taken as the promoter sequence, and Primerstar (Takara, Beijing) was used to clone the promoter. PlantPAN 3.0 (http://plantpan.itps.ncku.edu.tw) (Chow *et al*., 2019) was used to predict the position of the NAC element on the promoter. Primers are shown in Table S1.

### Yeast two-hybrid (Y1H) assay

The full-length coding sequence *SmNAC* was cloned into pGADT7. The promoter fragment of *SmDDA1b* was ligated into the pAbAi vector to generate baits. The Y1H experiment was carried out according to the manufacturer’s protocol for the Matchmaker Gold Y1H library screening system (Clontech, USA). The primers are listed in Table S1.

### Dual-luciferase assay

The 1474 bp promoters of *SmDDA1b* was inserted into the pGreen II 0800-LUC vector as reporters, while pGreenII 62-SK-*SmNAC*, pGreenII 62-SK-*SmDDA1b* and empty pGreenII 62-SK served as effectors. The *Agrobacterium tumefaciens* strain GV3101 containing the corresponding effectors and reporters (v: v, 20: 1) were infiltrated into healthy *N. benthamiana* leaves. After incubation for 24-36 h, MG132 (50 μM) was injected into leaves. After incubation for three to four days, the firefly LUC and Renilla LUC activities were measured by Dual Luciferase Reporter Gene Assay Kit (Yeasen, Shanghai) and Cytation 5 Cell Imaging Multi-Mode Reader (BioTek, USA). Activity is expressed as the ratio of firefly LUC activity to Renilla LUC activity. The primers are listed in Table S1.

### Accession numbers

The GenBank accession number of *SmDDA1b*: MZ736671.

## Supporting Information

Fig. S1. SmDDA1b gene and amino acid sequence.

Fig. S2. SmDDA1b gene cDNA sequence in E31 and E32.

Fig. S3. Detection of disease resistance of E31 and E32 to *Ralstonia solanacearum* species complex (RSSC).

Fig. S4. The subcellular localization results of SmDDA1b.

Fig. S5. There is no interaction between SmDDA1b and ICS1.

Fig. S6. *SmDDA1b* promotor sequence.

Fig. S7. *SmDDA1bpro-2-3* has self-activation.

Table S1 List of primers.

Table S2 List of primers used for qPCR.

Table S3 A statistical table of 14 species and their genome accession numbers used in the *SmDDA1b* phylogenetic tree except for eggplant.

Table S4 List of plant incidence rate and disease index for testing the resistance of E31 and E32 to *Ralstonia solanacearum* species complex.

Table S5 List of *bar* gene test results of T_1_ generation.

Table S6 List of plant incidence rate and disease index in overexpression experiment.

Table S7 List of NAC transcription factor binding sites.

## Acknowledgements

We thank Lianhui Zhang (South China Agricultural University) for providing GMI1000 strain.

## Parsed Citations

Adams EHG, Spoel SH (2018) The ubiquitin–proteasome system as a transcriptional regulator of plant immunity. Journal of Experimental Botany 69: 4529–4537

Barchi L, Rabanus-Wallace MT, Prohens J, Toppino L, Padmarasu S, Portis E, Rotino GL, Stein N, Lanteri S, Giuliano G (2021) Improved genome assembly and pan-genome provide key insights on eggplant domestication and breeding. The Plant Journal

Boschi F, Schvartzman C, Murchio S, Ferreira V, Siri MI, Galván GA, Smoker M, Stransfeld L, Zipfel C, Vilaró FL (2017) Enhanced bacterial wilt resistance in potato through expression of Arabidopsis EFR and introgression of quantitative resistance from Solanum commersonii. Frontiers in plant science 8: 1642

Chang Y, Yu R, Feng J, Chen H, Eri H, Gao G (2020) NAC transcription factor involves in regulating bacterial wilt resistance in potato. Functional Plant Biology 47: 925–936

Checker VG, Kushwaha HR, Kumari P, Yadav S (2018) Role of phytohormones in plant defense: signaling and cross talk. In Molecular aspects of plant-pathogen interaction. Springer, pp 159–184

Chen C, Chen H, Zhang Y, Thomas HR, Frank MH, He Y, Xia R (2020) TBtools: an integrative toolkit developed for interactive analyses of big biological data. Molecular plant 13: 1194–1202

Chen IH, Chang JE, Wu CY, Huang YP, Hsu YH, Tsai CH (2019) An E3 ubiquitin ligase from Nicotiana benthamiana targets the replicase of Bamboo mosaic virus and restricts its replication. Molecular plant pathology 20: 673–684

Chen YY, Lin YM, Chao TC, Wang JF, Liu AC, Ho FI, Cheng CP (2009) Virus-induced gene silencing reveals the involvement of ethylene-, salicylic acid-and mitogen-activated protein kinase-related defense pathways in the resistance of tomato to bacterial wilt. Physiologia Plantarum 136: 324–335

Chen Z-Q, Zhao J-H, Chen Q, Zhang Z-H, Li J, Guo Z-X, Xie Q, Ding S-W, Guo H-S (2020) DNAgeminivirus infection induces an imprinted E3 ligase gene to epigenetically activate viral gene transcription. Plant Cell 32: 3256–3272

Chico JM, Lechner E, Fernandez-Barbero G, Canibano E, García-Casado G, Franco-Zorrilla JM, Hammann P, Zamarreño AM, García-Mina JM, Rubio V (2020) CUL3BPM E3 ubiquitin ligases regulate MYC2, MYC3, and MYC4 stability and JAresponses. Proceedings of the National Academy of Sciences 117: 6205–6215

Choi C, Hwang SH, Fang IR, Kwon SI, Park SR, Ahn I, Kim JB, Hwang DJ (2015) Molecular characterization of Oryza sativa WRKY 6, which binds to W-box-like element 1 of the Oryza sativa pathogenesis-related (PR) 10a promoter and confers reduced susceptibility to pathogens. New Phytologist 208: 846–859

Chow C-N, Lee T-Y, Hung Y-C, Li G-Z, Tseng K-C, Liu Y-H, Kuo P-L, Zheng H-Q, Chang W-C (2019) PlantPAN3. 0: a new and updated resource for reconstructing transcriptional regulatory networks from ChIP-seq experiments in plants. Nucleic acids research 47: D1155–D1163

Cui H, Gobbato E, Kracher B, Qiu J, Bautor J, Parker JE (2017) Acore function of EDS1 with PAD4 is to protect the salicylic acid defense sector in Arabidopsis immunity. New Phytologist 213: 1802–1817

Dempsey DMA, Vlot AC, Wildermuth MC, Klessig DF (2011) Salicylic acid biosynthesis and metabolism. The Arabidopsis book/American Society of Plant Biologists 9

Derksen H, Rampitsch C, Daayf F (2013) Signaling cross-talk in plant disease resistance. Plant science 207: 79–87

Ding M, Zhang K, He Y, Zuo Q, Zhao H, He M, Georgiev MI, Park SU, Zhou M (2021) FtBPM3 modulates the orchestration of FtMYB11-mediated flavonoids biosynthesis in Tartary buckwheat. Plant Biotechnology Journal 19: 1285

Ding Y, Sun T, Ao K, Peng Y, Zhang Y, Li X, Zhang Y (2018) Opposite roles of salicylic acid receptors NPR1 and NPR3/NPR4 in transcriptional regulation of plant immunity. Cell 173: 1454-1467. e1415

Drugeon G, Jupin I (2002) Stability in vitro of the 69K movement protein of Turnip yellow mosaic virus is regulated by the ubiquitin-mediated proteasome pathway. Journal of general virology 83: 3187–3197

Eddy SR (1998) Profile hidden Markov models. Bioinformatics (Oxford, England) 14: 755–763

Feys BJ, Moisan LJ, Newman MA, Parker JE (2001) Direct interaction between the Arabidopsis disease resistance signaling proteins, EDS1 and PAD4. The EMBO journal 20: 5400–5411

Garcion C, Lohmann A, Lamodière E, Catinot J, Buchala A, Doermann P, Métraux J-P (2008) Characterization and biological function of the ISOCHORISMATE SYNTHASE2 gene of Arabidopsis. Plant physiology 147: 1279–1287

Gong C, Su H, Li Z, Mai P, Sun B, Li Z, Heng Z, Xu X, Yang S, Li T (2021) Involvement of histone acetylation in tomato resistance to Ralstonia solanacearum. Scientia Horticulturae 285: 110163

Hayward A(1991) Biology and epidemiology of bacterial wilt caused by Pseudomonas solanacearum. Annual review of phytopathology 29: 65–87

Hernández Y, Sanan-Mishra N (2017) miRNA mediated regulation of NAC transcription factors in plant development and environment stress response. Plant Gene 11: 190–198

Huang Q, Allen C (2000) Polygalacturonases are required for rapid colonization and full virulence of Ralstonia solanacearum on tomato plants. Physiological and molecular plant pathology 57: 77–83

Hussain A, Li X, Weng Y, Liu Z, Ashraf MF, Noman A, Yang S, Ifnan M, Qiu S, Yang Y (2018) CaWRKY22 acts as a positive regulator in pepper response to Ralstonia solanacearum by constituting networks with CaWRKY6, CaWRKY27, CaWRKY40, and CaWRKY58. International journal of molecular sciences 19: 1426

Irigoyen ML, Iniesto E, Rodriguez L, Puga MI, Yanagawa Y, Pick E, Strickland E, Paz-Ares J, Wei N, De Jaeger G (2014) Targeted degradation of abscisic acid receptors is mediated by the ubiquitin ligase substrate adaptor DDA1 in Arabidopsis. The Plant Cell 26: 712–728

Jia C, Zhang L, Liu L, Wang J, Li C, Wang Q (2013) Multiple phytohormone signalling pathways modulate susceptibility of tomato plants to Alternaria alternata f. sp. lycopersici. Journal of experimental botany 64: 637–650

Karki SJ, Reilly A, Zhou B, Mascarello M, Burke J, Doohan F, Douchkov D, Schweizer P, Feechan A(2021) Asmall secreted protein from Zymoseptoria tritici interacts with a wheat E3 ubiquitin ligase to promote disease. Journal of experimental botany 72: 733–746

Katoh K, Standley DM (2013) MAFFT multiple sequence alignment software version 7: improvements in performance and usability. Molecular biology and evolution 30: 772–780

Kim B-S, French E, Caldwell D, Harrington EJ, Iyer-Pascuzzi AS (2016) Bacterial wilt disease: Host resistance and pathogen virulence mechanisms. Physiological and Molecular Plant Pathology 95: 37–43

Kunwar S, Iriarte F, Fan Q, Evaristo da Silva E, Ritchie L, Nguyen NS, Freeman JH, Stall RE, Jones JB, Minsavage GV (2018) Transgenic expression of EFR and Bs2 genes for field management of bacterial wilt and bacterial spot of tomato. Phytopathology 108: 1402–1411

Lee D, Lal NK, Lin Z-JD, Ma S, Liu J, Castro B, Toruño T, Dinesh-Kumar SP, Coaker G (2020) Regulation of reactive oxygen species during plant immunity through phosphorylation and ubiquitination of RBOHD. Nature communications 11: 1–16

Lemessa F, Zeller W (2007) Isolation and characterisation of Ralstonia solanacearum strains from Solanaceae crops in Ethiopia. Journal of Basic Microbiology 47: 40–49

Li F, Huang C, Li Z, Zhou X (2014) Suppression of RNAsilencing by a plant DNAvirus satellite requires a host calmodulin-like protein to repress RDR6 expression. PLoS pathogens 10: e1003921

Li N, Han X, Feng D, Yuan D, Huang L-J (2019) Signaling crosstalk between salicylic acid and ethylene/jasmonate in plant defense: do we understand what they are whispering? International Journal of Molecular Sciences 20: 671

Li S, Lin Y-CJ, Wang P, Zhang B, Li M, Chen S, Shi R, Tunlaya-Anukit S, Liu X, Wang Z (2019) The AREB1 transcription factor influences histone acetylation to regulate drought responses and tolerance in Populus trichocarpa. The Plant Cell 31: 663–686

Lindgren PB (1997) The role of hrp genes during plant-bacterial interactions. Annual review of phytopathology 35: 129–152

Liu Q, Liu Y, Tang Y, Chen J, Ding W (2017) Overexpression of NtWRKY50 increases resistance to Ralstonia solanacearum and alters salicylic acid and jasmonic acid production in tobacco. Frontiers in plant science 8: 1710

Liu Y, Hou J, Wang X, Li T, Majeed U, Hao C, Zhang X (2020) The NAC transcription factor NAC019-A1 is a negative regulator of starch synthesis in wheat developing endosperm. Journal of experimental botany 71: 5794–5807

Liu Y, Tang Y, Tan X, Ding W (2021) NtRNF217, Encoding a Putative RBR E3 Ligase Protein of Nicotiana tabacum, Plays an Important Role in the Regulation of Resistance to Ralstonia solanacearum Infection. International journal of molecular sciences 22: 5507

Livak KJ, Schmittgen TD (2001) Analysis of relative gene expression data using real-time quantitative PCR and the 2− ΔΔCT method. methods 25: 402–408

Lowe-Power TM, Jacobs JM, Ailloud F, Fochs B, Prior P, Allen C (2016) Degradation of the plant defense signal salicylic acid protects Ralstonia solanacearum from toxicity and enhances virulence on tobacco. MBio 7: e00656–00616

Lu P-P, Yu T-F, Zheng W-J, Chen M, Zhou Y-B, Chen J, Ma Y-Z, Xi Y-J, Xu Z-S (2018) The wheat Bax Inhibitor-1 protein interacts with an aquaporin TaPIP1 and enhances disease resistance in Arabidopsis. Frontiers in plant science 9: 20

Ma J, Chen J, Wang M, Ren Y, Wang S, Lei C, Cheng Z (2018) Disruption of OsSEC3Aincreases the content of salicylic acid and induces plant defense responses in rice. Journal of experimental botany 69: 1051–1064

Mahesh HM, Sharada MS (2018) Histopathological response of resistance induced by salicylic acid during brinjal (Solanum melongena L.) -Verticillium dahliae interaction. Journal of Applied Biology & Biotechnology 6

Mansfield J, Genin S, Magori S, Citovsky V, Sriariyanum M, Ronald P, Dow M, Verdier V, Beer SV, Machado MA(2012) Top 10 plant pathogenic bacteria in molecular plant pathology. Molecular plant pathology 13: 614–629

Marino D, Froidure S, Canonne J, Khaled SB, Khafif M, Pouzet C, Jauneau A, Roby D, Rivas S (2013) Arabidopsis ubiquitin ligase MIEL1 mediates degradation of the transcription factor MYB30 weakening plant defence. Nature communications 4: 1–9

Marques ANJ, Palanimurugan R, Matias AC, Ramos PC, Dohmen RJR (2009) Catalytic mechanism and assembly of the proteasome. Chemical Reviews 109: 1509–1536

McGarvey J, Denny T, Schell M (1999) Spatial-temporal and quantitative analysis of growth and EPS I production by Ralstonia solanacearum in resistant and susceptible tomato cultivars. Phytopathology 89: 1233–1239

McLellan H, Chen K, He Q, Wu X, Boevink PC, Tian Z, Birch PR (2020) The ubiquitin E3 ligase PUB17 positively regulates immunity by targeting a negative regulator, KH17, for degradation. Plant communications 1: 100020

Miao M, Niu X, Kud J, Du X, Avila J, Devarenne TP, Kuhl JC, Liu Y, Xiao F (2016) The ubiquitin ligase SEVEN IN ABSENTIA (SINA) ubiquitinates a defense-related NAC transcription factor and is involved in defense signaling. New Phytol 211: 138–148

Morel A, Guinard J, Lonjon F, Sujeeun L, Barberis P, Genin S, Vailleau F, Daunay MC, Dintinger J, Poussier S (2018) The eggplant AG91-25 recognizes the type III-secreted effector RipAX2 to trigger resistance to bacterial wilt (Ralstonia solanacearum species complex). Molecular plant pathology 19: 2459–2472

Morreale FE, Walden H (2016) Types of Ubiquitin Ligases. Cell 165: 248–248 e241

Na C, Shuanghua W, Jinglong F, Bihao C, Jianjun L, Changming C, Jin J (2016) Overexpression of the eggplant (Solanum melongena) NAC family transcription factor S mNAC suppresses resistance to bacterial wilt. Scientific reports 6: 1–20

Nakano M, Mukaihara T (2018) Ralstonia solanacearum type III effector RipAL targets chloroplasts and induces jasmonic acid production to suppress salicylic acid-mediated defense responses in plants. Plant and Cell Physiology 59: 2576–2589

Nakano M, Oda K, Mukaihara T (2017) Ralstonia solanacearum novel E3 ubiquitin ligase (NEL) effectors RipAW and RipAR suppress pattern-triggered immunity in plants. Microbiology 163: 992–1002

Narusaka M, Hatakeyama K, Shirasu K, Narusaka Y (2014) Arabidopsis dual resistance proteins, both RPS4 and RRS1, are required for resistance to bacterial wilt in transgenic Brassica crops. Plant signaling & behavior 9: e29130

Nguyen L-T, Schmidt HA, Von Haeseler A, Minh BQ (2015) IQ-TREE: a fast and effective stochastic algorithm for estimating maximum-likelihood phylogenies. Molecular biology and evolution 32: 268–274

O’Leary NA, Wright MW, Brister JR, Ciufo S, Haddad D, McVeigh R, Rajput B, Robbertse B, Smith-White B, Ako-Adjei D (2016) Reference sequence (RefSeq) database at NCBI: current status, taxonomic expansion, and functional annotation. Nucleic acids research 44: D733–D745

Oh S-K, Kim H, Choi D (2014) Rpi-blb2-mediated late blight resistance in Nicotiana benthamiana requires SGT1 and salicylic acid-mediated signaling but not RAR1 or HSP90. FEBS letters 588: 1109–1115

Olma MH, Roy M, Le Bihan T, Sumara I, Maerki S, Larsen B, Quadroni M, Peter M, Tyers M, Pintard L (2009) An interaction network of the mammalian COP9 signalosome identifies Dda1 as a core subunit of multiple Cul4-based E3 ligases. Journal of cell science 122: 1035–1044

Pang P-X, Shi L, Wang X-J, Chang Y-N, Luo Y-P, Feng J-L, Eri H, Gao G (2019) Cloning and expression analysis of the StCUL1 gene in potato. Journal of Plant Biochemistry and Biotechnology 28: 460–469

Park S-R, Cha E-M, Kim T-H, Han S-Y, Hwang D-J, Ahn I-P, Cho K-S, Bae S-C (2012) Isolation of Potato StACRE Gene and Its Function in Resistance against Bacterial Wilt Disease. Journal of Life Science 22: 177–183

Patil VU, Gopal J, Singh B (2012) Improvement for bacterial wilt resistance in potato by conventional and biotechnological approaches. Agricultural research 1: 299–316

Pauwels L, Ritter A, Goossens J, Durand AN, Liu H, Gu Y, Geerinck J, Boter M, Vanden Bossche R, De Clercq R (2015) The ring e3 ligase keep on going modulates jasmonate zim-domain12 stability. Plant Physiology 169: 1405–1417

Pickart CM, Fushman D (2004) Polyubiquitin chains: polymeric protein signals. Curr Opin Chem Biol 8: 610–616

Poueymiro M, Genin S (2009) Secreted proteins from Ralstonia solanacearum: a hundred tricks to kill a plant. Current opinion in microbiology 12: 44–52

Qiu Z, Wang X, Gao J, Guo Y, Huang Z, Du Y (2016) The tomato Hoffman’s anthocyaninless gene encodes a bHLH transcription factor involved in anthocyanin biosynthesis that is developmentally regulated and induced by low temperatures. PloS one 11: e0151067

Qiu Z, Yan S, Xia B, Jiang J, Yu B, Lei J, Chen C, Chen L, Yang Y, Wang Y (2019) The eggplant transcription factor MYB44 enhances resistance to bacterial wilt by activating the expression of spermidine synthase. Journal of experimental botany 70: 5343–5354

Rowland O, Ludwig AA, Merrick CJ, Baillieul F, Tracy FE, Durrant WE, Fritz-Laylin L, Nekrasov V, Sjolander K, Yoshioka H, Jones JD (2005) Functional analysis of Avr9/Cf-9 rapidly elicited genes identifies a protein kinase, ACIK1, that is essential for full Cf-9-dependent disease resistance in tomato. Plant Cell 17: 295–310

Safni I, Cleenwerck I, De Vos P, Fegan M, Sly L, Kappler U (2014) Polyphasic taxonomic revision of the Ralstonia solanacearum species complex: proposal to emend the descriptions of Ralstonia solanacearum and Ralstonia syzygii and reclassify current R. syzygii strains

as Ralstonia syzygii subsp. syzygii subsp. nov., R. solanacearum phylotype IV strains as Ralstonia syzygii subsp. indonesiensis subsp. nov., banana blood disease bacterium strains as Ralstonia syzygii subsp. celebesensis subsp. nov. and R. solanacearum phylotype I and III strains as Ralstoniapseudosolanacearum sp. nov. International journal of systematic and evolutionary microbiology 64: 3087–3103

Shabek N, Ruble J, Waston CJ, Garbutt KC, Hinds TR, Li T, Zheng N (2018) Structural insights into DDA1 function as a core component of the CRL4-DDB1 ubiquitin ligase. Cell discovery 4: 1–4

Shen Q, Hu T, Bao M, Cao L, Zhang H, Song F, Xie Q, Zhou X (2016) Tobacco RING E3 ligase NtRFP1 mediates ubiquitination and proteasomal degradation of a geminivirus-encoded βC1. Molecular plant 9: 911–925

Stone SL, Hauksdóttir H, Troy A, Herschleb J, Kraft E, Callis J (2005) Functional Analysis of the RING-Type Ubiquitin Ligase Family of Arabidopsi s. Plant physiology 137: 13–30

Sun B, Zhou X, Chen C, Chen C, Chen K, Chen M, Liu S, Chen G, Cao B, Cao F (2020) Coexpression network analysis reveals an MYB transcriptional activator involved in capsaicinoid biosynthesis in hot peppers. Horticulture research 7: 1–14

Tahir HAS, Gu Q, Wu H, Niu Y, Huo R, Gao X (2017) Bacillus volatiles adversely affect the physiology and ultra-structure of Ralstonia solanacearum and induce systemic resistance in tobacco against bacterial wilt. Scientific reports 7: 1–15

Tasset C, Bernoux M, Jauneau A, Pouzet C, Brière C, Kieffer-Jacquinod S, Rivas S, Marco Y, Deslandes L (2010) Autoacetylation of the Ralstonia solanacearum effector PopP2 targets a lysine residue essential for RRS1-R-mediated immunity in Arabidopsis. PLoS pathogens 6: e1001202

Thrower JS, Hoffman L, Rechsteiner M, Pickart CM (2000) Recognition of the polyubiquitin proteolytic signal. The EMBO journal 19: 94–102

Tong S, Chen N, Wang D, Ai F, Liu B, Ren L, Chen Y, Zhang J, Lou S, Liu H (2021) The U-box E3 ubiquitin ligase PalPUB79 positively regulates ABA-dependent drought tolerance via ubiquitination of PalWRKY77 in Populus. Plant Biotechnology Journal

Wang H, Cheng Z, Wang B, Dong J, Ye W, Yu Y, Liu T, Cai X, Song B, Liu J (2020) Combining genome composition and differential gene expression analyses reveals that SmPGH1 contributes to bacterial wilt resistance in somatic hybrids. Plant Cell Reports 39: 1235–1248

Wang W, Fan Y, Niu X, Miao M, Kud J, Zhou B, Zeng L, Liu Y, Xiao F (2018) Functional analysis of the seven in absentia ubiquitin ligase family in tomato. Plant, cell & environment 41: 689–703

Wang Y, Yu B, Yan S, Qiu Z, Chen C, Lei J, Tian S, Cao B (2020) Advances in research on plant ubiquitin genes. Chinese Agricultural Science Bulletin 36: 14–22

Wiermer M, Feys BJ, Parker JE (2005) Plant immunity: the EDS1 regulatory node. Current opinion in plant biology 8: 383–389

Wildermuth MC, Dewdney J, Wu G, Ausubel FM (2001) Isochorismate synthase is required to synthesize salicylic acid for plant defence. Nature 414: 562–565

Wu Y, Zhang D, Chu JY, Boyle P, Wang Y, Brindle ID, De Luca V, Després C (2012) The Arabidopsis NPR1 protein is a receptor for the plant defense hormone salicylic acid. Cell reports 1: 639–647

Xi-ou X, Jing J, Na C, Jian-jun L, Ling-ling L, Bi-hao C (2016) Identification of Key Signal Gene Involved in Eggplant Bacterial Wilt-resistance. Acta Horticulturae Sinica 43: 1295

Xi’ou X, Bihao C, Guannan L, Jianjun L, Qinghua C, Jin J, Yujing C (2015) Functional characterization of a putative bacterial wilt resistance gene (RE-bw) in eggplant. Plant molecular biology reporter 33: 1058–1073

Xie Q, Guo H-S, Dallman G, Fang S, Weissman AM, Chua N-H (2002) SINAT5 promotes ubiquitin-related degradation of NAC1 to attenuate auxin signals. Nature 419: 167–170

Yanagawa Y, Sullivan JA, Komatsu S, Gusmaroli G, Suzuki G, Yin J, Ishibashi T, Saijo Y, Rubio V, Kimura S (2004) Arabidopsis COP10 forms a complex with DDB1 and DET1 in vivo and enhances the activity of ubiquitin conjugating enzymes. Genes & development 18: 2172–2181

Yoshii M, Yamazaki M, Rakwal R, Kishi-Kaboshi M, Miyao A, Hirochika H (2010) The NAC transcription factor RIM1 of rice is a new regulator of jasmonate signaling. The Plant Journal 61: 804–815

Yu G, Xian L, Xue H, Yu W, Rufian JS, Sang Y, Morcillo RJ, Wang Y, Macho AP (2020) Abacterial effector protein prevents MAPK-mediated phosphorylation of SGT1 to suppress plant immunity. PLoS pathogens 16: e1008933

Zhang C, Wei Y, Xu L, Wu K-C, Yang L, Shi C-N, Yang G-Y, Chen D, Yu F-F, Xie Q (2020) ABunyavirus-inducible ubiquitin ligase targets RNApolymerase IV for degradation during viral pathogenesis in rice. Molecular plant 13: 836–850

Zhang L, Yao L, Zhang N, Yang J, Zhu X, Tang X, Calderón-Urrea A, Si H (2018) Lateral root development in potato is mediated by stu-mi164 regulation of NAC transcription factor. Frontiers in plant science 9: 383

Zhang Z, Chen H, Huang X, Xia R, Zhao Q, Lai J, Teng K, Li Y, Liang L, Du Q (2011) BSCTV C2 attenuates the degradation of SAMDC1 to suppress DNA methylation-mediated gene silencing in Arabidopsis. The Plant Cell 23: 273–288

Zhao XY, Qi CH, Jiang H, Zhong MS, You CX, Li YY, Hao YJ (2020) MdWRKY15 improves resistance of apple to Botryosphaeria dothidea via the salicylic acid-mediated pathway by directly binding the MdICS1 promoter. Journal of integrative plant biology 62: 527–543

Zhu Y, Yan J, Liu W, Liu L, Sheng Y, Sun Y, Li Y, Scheller HV, Jiang M, Hou X (2016) Phosphorylation of a NAC transcription factor by a calcium/calmodulin-dependent protein kinase regulates abscisic acid-induced antioxidant defense in maize. Plant Physiology 171: 1651–1664

Zimmerman ES, Schulman BA, Zheng N (2010) Structural assembly of cullin-RING ubiquitin ligase complexes. Current opinion in structural biology 20: 714–721

